# Molecular Mechanism of pH Sensing and Activation in GPR4 Reveals Proton-Mediated GPCR Signaling

**DOI:** 10.1101/2025.02.16.638498

**Authors:** Chongzhao You, Shimeng Guo, Tianwei Zhang, Xinheng He, Tianyu Gao, Wenwen Xin, Zining Zhu, Yujie Lu, Youwei Xu, Zhen Li, Yumu Zhang, Xi Cheng, Yi Jiang, Xin Xie, H. Eric Xu

## Abstract

Maintaining pH homeostasis is critical for cellular function across all living organisms. Proton-sensing G protein-coupled receptors (GPCRs), particularly GPR4, play a pivotal role in cellular responses to pH changes, yet the molecular mechanisms underlying their proton sensing and activation remain incompletely understood. Here we present high-resolution cryo-electron microscopy structures of GPR4 in complex with G proteins under physiological and acidic pH conditions. Our structures expose an intricate proton-sensing mechanism driven by a sophisticated histidine network in the receptor’s extracellular domain. Upon protonation of key histidines under acidic conditions, a remarkable conformational cascade is initiated, propagating from the extracellular region to the intracellular G protein-coupling interface. This dynamic process involves precise transmembrane helix rearrangements and conformational shifts of conserved motifs, mediated by strategically positioned water molecules. Notably, we discovered a bound bioactive lipid, lysophosphatidylcholine, which has positive allosteric effects on GPR4 activation. These findings provide a comprehensive framework for understanding proton sensing in GPCRs and the interplay between pH sensing and lipid regulation, offering insights into cellular pH homeostasis and potential therapies for pH-related disorders.

## Introduction

Maintaining physiological pH homeostasis within the narrow range of 7.35-7.45, is fundamental to cellular function and organismal survival ^1–4^. Disruption of this delicate balance occurs in numerous pathological conditions, including diabetes, renal dysfunction, respiratory disorders, and cancer, as well as during intense physical activity and dietary changes ^2,5–10^. The human body employs sophisticated regulatory mechanisms to maintain pH homeostasis, encompassing buffer systems, respiratory and renal regulation, and cellular ion exchange ^1,2,5,11^. At the molecular level, three distinct classes of pH sensors orchestrate cellular responses: proton-sensing G protein-coupled receptors (GPCRs), proton transporters (H+-ATPases), and acid-sensing ion channels (ASICs) ^12–16^. While the structures and activation mechanisms of H^+^-ATPases and ASICs have been extensively characterized structurally and mechanistically ^16–18^, the molecular basis of proton sensing by GPCRs remains unclear.

GPR4, a prominent member of proton-sensing GPCRs, commands particular attention due to its ubiquitous expression and crucial roles in endothelial function, tumor biology, and metabolic acidosis regulation ^7,19–24^. GPR4, a prominent member of proton-sensing GPCRs, commands particular attention due to its ubiquitous expression and crucial roles in endothelial function, tumor biology, and metabolic acidosis regulation, maintaining partial activity at physiological pH while achieving full activation under acidic conditions ^25,26^. Although GPR4 predominantly signals through G_s_ proteins, it exhibits remarkable coupling plasticity across G protein subtypes. Notably, the G_q_ pathway demonstrates a distinct pH-sensing range, potentially suggesting an alternative sensing mechanism ^27^. Two popular theories have emerged to explain the proton-sensing mechanism of GPR4. One is centered on extracellular histidine residues ^25,28–30^, and another focuses on pK_a_ shifts in buried acidic residues ^26,31,32^. Recent evolutionary analysis has highlighted the critical role of the extracellular domain (ECD) in proton sensing ^33^, while functional studies of related receptors, particularly GPR68, have provided valuable insights into pH sensitivity mechanisms ^34^. However, the precise molecular mechanism of human GPR4 proton sensing remains to be fully elucidated.

A fascinating aspect of GPR4 regulation involves lysophosphatidylcholine (LPC), an abundant plasma membrane lipid ^35,36^. LPC is associated with endothelial functions and immune regulation, while serving as a precursor for bioactive lipids through autotaxin-mediated conversion ^36–38^. Evidence suggests LPC modulates GPR4 activity and potentially mediates both lysophospholipid-dependent and -independent pathways in tumor development ^35,36,38^. However, the molecular basis of LPC-GPR4 interactions and their physiological significance remains poorly understood.

The clinical relevance of GPR4 stems from its pivotal role in endothelial cell regulation and its implications in various pathological conditions ^7,19,20,39–41^. Under acidic conditions, GPR4 activation promotes angiogenesis and enhances vascular permeability, potentially facilitating tumor growth and metastasis while perpetuating acidosis ^10,19,23,39,42^. These findings have established GPR4 as a promising therapeutic target, spurring efforts to develop selective antagonists ^43–46^.

To address the critical need for mechanistic understanding in drug development, we present high-resolution structures of GPR4 in complex with G_s_ and G_q_ transducers under physiological (pH 7.4) and acidic (pH 6.5) conditions. Our structural analysis reveals critical insights into proton sensing and activation mechanisms. Furthermore, we identify and characterize LPC binding sites, illuminating their functional impact on GPR4 activation. Through comprehensive structural analysis, pharmacological profiling, mutagenesis, and computational approaches, we provide a detailed molecular framework of GPR4’s proton-sensing mechanism and lipid interactions. These findings not only advance our knowledge of pH-sensing GPCRs but also establish a foundation for structure-based drug design targeting pH-related pathologies.

## Results

### Unique conformations of GPR4 complexes

GPR4 exhibits a distinct pH-dependent activation profile, responding to pH changes between 5.8 and 7.8, with inactivation occurring above pH 7.8, as confirmed by cAMP accumulation assays ^27^ (Fig. 1a-b). To elucidate the molecular mechanisms of proton sensing and activation in GPR4, we determined high-resolution cryo-EM structures of GPR4 in complex with G_s_ under both physiological (pH 7.4) and acidic (pH 6.5) conditions at resolutions of 2.59 Å and 2.36 Å, respectively. We also determined the cryo-EM structure of GPR4 in complex with G_q_ proteins at a resolution of 2.55 Å at pH 7.4 (Fig. 1d-f and Fig. S1-3). Our density maps enabled precise modeling of the receptor, G proteins, and associated components (Fig. S4).

**Fig. 1.**
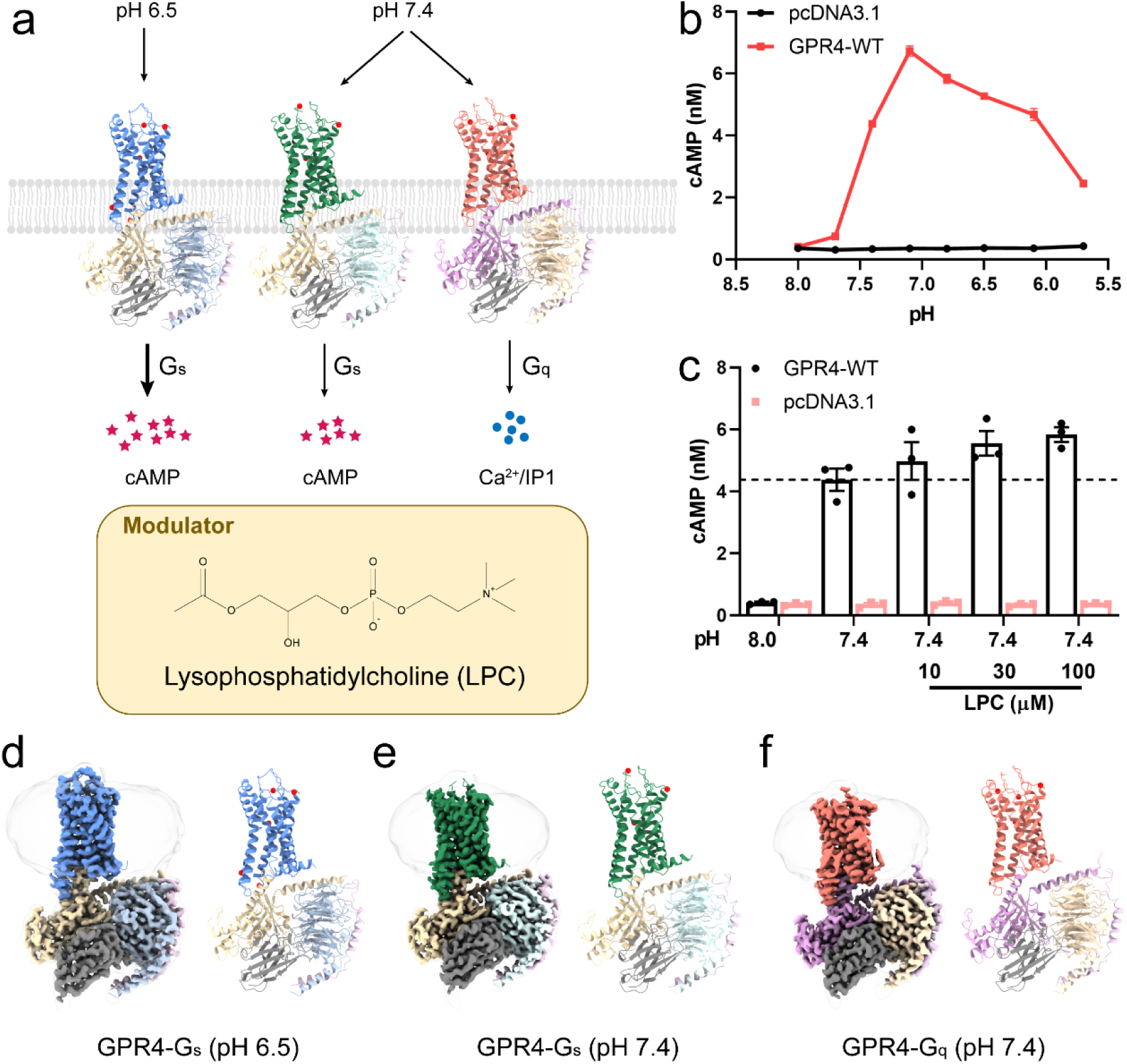
Overall structures of GPR4-G_s_/G_q_ complexes at acidic and physiological pH. **a** Schema diagram for GPR4 at various pH activating G_s_/G_q_ pathway. **b** Curves showing pH-dependent cAMP accumulation in cells overexpressing GPR4 or pcDNA3.1. **c** LPC induced activation of GPR4 at pH 7.4 displayed as dose-dependent measurement by cAMP accumulation assay. Values are represented as mean ± SEM of three independent experiments (n = 3). **d-f** Overall structures and EM-density map for GPR4 complexes. Colors are shown as indicated. Red balls refer to water molecules in our structures.

A defining feature of active GPR4 structures is the cooperative organization of ECD. Specifically, N-terminus and TM1 exhibit a distinctive bend toward the central region of the receptor, while extracellular loops (ECLs) form stabilizing interactions with N-terminus through an intricate network of water molecules (Figure 2a). Surface analysis reveals a highly acidic and hydrophilic ECD, a characteristic crucial for proton sensing^34^ (Fig. 2b-c). The high resolution of our structures (2.36-2.59 Å) allows precise mapping of these water molecules, providing important insights into their role in proton sensing and signal (Fig. S4d).

**Fig. 2.**
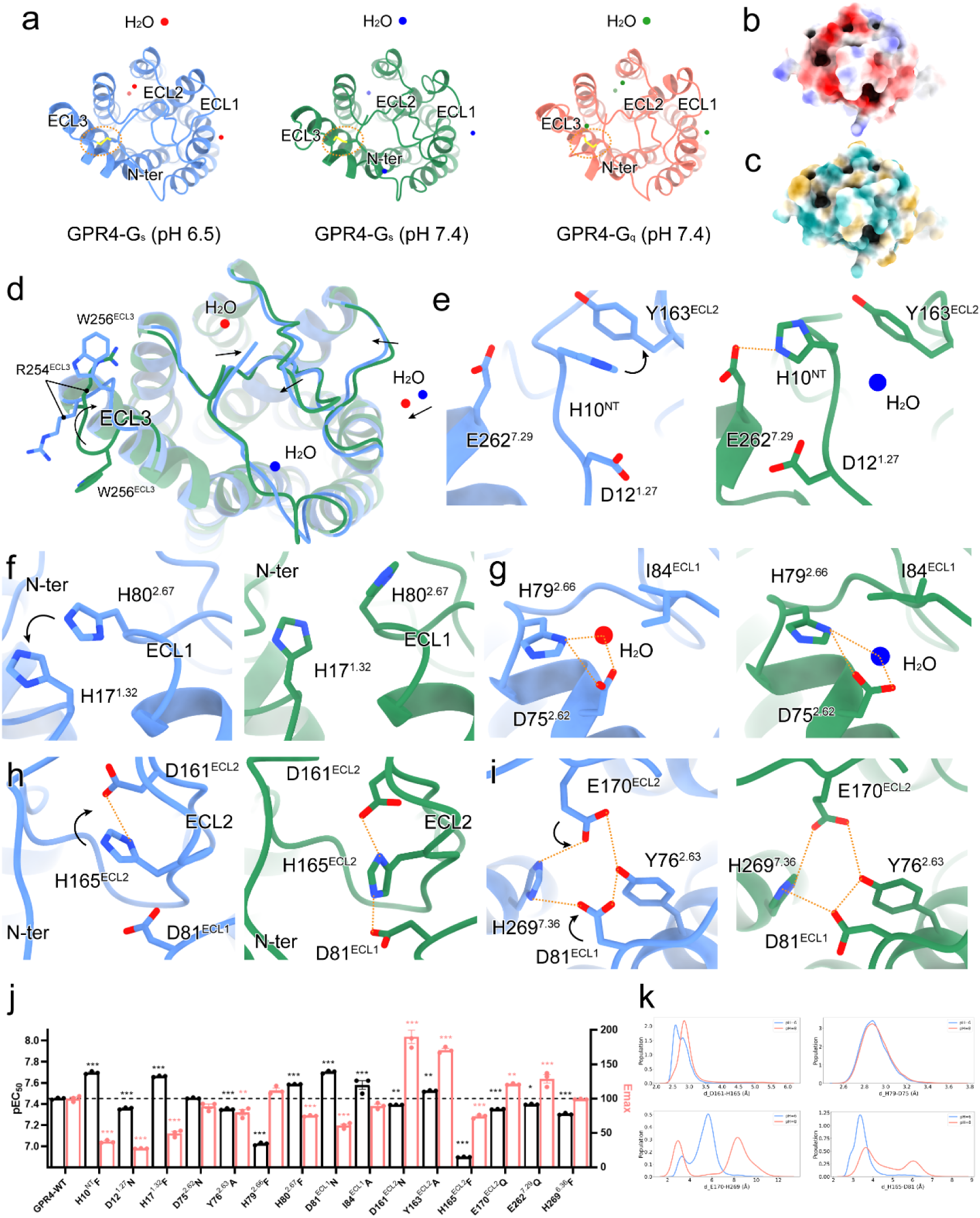
Proton recognition mode of GPR4. **a** The top view of active GPR4 with the distribution of protons around ECD. The disulfide bonds are labeled by orange dashed circles and the water molecules are displayed as sphere. Colors are shown as indicated. **b-c** The surface of ECD in the GPR4-G_s_ complex at pH 6.5 by electrostatic (b) and hydrophobic (c) analyses. **d** The structural superposition of ECD in the GPR4-G_s_ complexes at pH 6.5 and 7.4. **e-i** Detailed interactions and comparisons of GPR4 at pH arrows and orange dashed lines, respectively. **j** Effects of mutations of GPR4 on the potency of pH-induced cAMP accumulation. The black bars represent the pEC_50_ values of pH-induced responses in GPR4 (WT) and mutants, while the red bars indicate the maximum cAMP concentrations induced by pH in GPR4 (WT) and mutants, both normalized to WT. A decrease in pEC_50_ indicates reduced sensitivity to pH. The original data are provided in Supplementary Fig. S9. Values are shown as mean ± S.E.M. from three independent experiments. **P<0.001 and ***P<0.0001 by one-way ANOVA followed by multiple comparisons test, compared with WT. **k** The extracellular conformation distribution under pH 6.0 and pH 8.0. The sidechain minimal distance distribution of D161^ECL2^-H165^ECL2^, D75^2.62^-H79^2.66^, E170^ECL2^-H269^7.36^, and D81^ECL1^-H269^7.36^.

Our structures reveal several unique architectural features that distinguish GPR4 from typical class A GPCRs. First, a rare disulfide bond between C9^NT^ and C258^7.25^, also present in GPR68 ^34^, appears to be a specialized feature of proton-sensing GPCRs (Fig. 2a and Fig. S5f). The bent TM1 of GPR4 forms extensive and compact networks with extracellular loops (ECLs) (Fig. 2a and Fig. S5). Mutation of either cysteine significantly impairs both proton sensitivity and receptor activation, highlighting the critical role of this structural element in receptor function ^34^ (Fig. S6). Second, the bent TM1 forms extensive networks with ECLs, creating a compact structure essential for proton sensing (Fig. 2a and Fig. S5). The structures exhibit hallmark features of GPCR activation, including pronounced outward movement of TM6 at the cytoplasmic end (Fig. S5a). However, GPR4 displays distinctive characteristics shared with other pH-sensing receptors like GPR65 and GPR68 ^34^ (Fig. S5f), including a unique two helical-turn extension of TM7 at the extracellular side, enabling ECL3-N-terminus interaction; inward movement of both N-terminus and the extracellular end of TM1, reminiscent of lipid-liganded GPCRs like GPR3, GPR40, and GPR119 ^47–50^; and an exceptionally compact ECD architecture where the N-terminus coordinates ECL1-3 interactions, crucial for proton sensing ^33,34^ (Fig. S5b-e).

Notably, our density maps revealed multiple associated lipids, including cholesterol and phospholipids (Fig. S4). Of particular interest, we identified clear density corresponding to lysophosphatidylcholine (LPC), previously proposed as an endogenous GPR4 ligand. Functional validation through cAMP accumulation assays demonstrated that LPC acts as a positive allosteric modulator of GPR4 in a dose-dependent manner (Fig. 1c). This finding aligns with recent reports of LPC ‘s role in modulating ADGRF1 ^51^ and GPR119 ^47^, suggesting a broader significance of LPC in GPCR signaling.

### The proton recognition mechanism of GPR4 complexes

Among proton-sensing GPCRs, GPR4 stands out due to its uniquely high histidine content (Fig. S7) and distinctive ability to maintain activation at elevated pH ranges (7.4 to 7.8) ^30^. The imidazole side chain of histidine serves as a precise proton receptor through its pH-dependent charge distribution ^52–54^, while aspartic and glutamic acid residues contribute to proton sensing through electrostatic interactions ^33,34,55,56^. The enrichment of both histidine and acidic amino acids in ECD of proton-sensing GPCRs suggests their orchestrated role in proton detection (Fig. S7).

Our high-quality reveal an extended polar network within ECD, comprising numerous proton-titratable residues, particularly in N-terminus and ECL2. Water molecules are clearly visible throughout the GPR4 ECD (Fig. 2a-c). While most of the ECD maintains similar conformations at pH 6.5 and 7.4, ECL3 exhibits distinct rotational changes (Fig. 2d). The complex at lower pH demonstrates tighter ECD association, accompanied by rotational shifts in R254^ECL3^ and W256^ECL3^ (Fig. 2d).

Comparative analysis of G_s_-coupled GPR4 at pH 6.5 (fully activated) and pH 7.4 (partially activated) reveals key conformational changes underlying proton sensing (Fig. 1b). At pH 6.5, the protonated H10^NT^ rotates toward Y163^ECL2^ to form a π-π interaction, while at pH 7.4, it interacts with E262^7.29^ and a water molecule that restrains its movement (Fig. 2e). The protonated H17^1.27^ and H80^2.67^ establish stable a stable π-π interaction at pH 6.5, maintaining N-terminus-ECL1 connectivity (Fig. 2f). A conserved water molecule occupies the cavity formed by D75^2.62^, H79^2.66^, and I84^ECL1^ at both pH conditions, creating a local polar network that stabilizes ECL1 conformation (Fig. 2g). The water molecule shifts slightly inward with H79^2.66^ rotation, triggering conformational changes (Fig. 2d, g). H165^ECL2^ undergoes inward rotation to coordinate with D161^ECL2^, forming strong electrostatic interactions (Fig. 2h), Additionally, Y76^2.63^, D81^ECL1^, and E170^ECL2^ generate a polar network that coordinates with the protonated H269^7.36^ (Fig. 2i). The evolutionarily conserved π-π stacking between H155^4.63^ and W177^5.34^ in non-human GPR4 ^33^ is preserved in our structures, with mutations disrupting this interaction significantly reducing receptor activity (Fig. S8). The pH-dependent conformational changes highlight the crucial role of these residues in proton sensing (Fig. 2j and Fig. S9). At pH 7.4, GPR4 maintains certain features, including interactions stabilizing N-terminus and ECLs conformation (Fig. 2d-i). Structural based pK_a_ calculations identify H269^7.36^ as the primary protonated histidine at pH 6.5 (Fig. S10), supported by molecular dynamics simulations analyzing specific residue sidechain minimal distance distributions (Fig. 2k).

The G_q_-coupled GPR4 at pH 7.4 shares conformational similarities with G_s_-coupled GPR4 complexes (Fig. 2a) while exhibiting distinct features. The H17^1.27^-H80^2.67^ π-π interaction and H79^2.66^-I84^ECL1^ water molecule network preserved (Fig. 3a-b). However, the H165^ECL2^-H269^7.36^ polar network shows expansion (Fig. 3c). D81^ECL1^ adopts a unique conformation, interacting with H80^2.67^ and H165^ECL2^ to create an enhanced polar environment (Fig. 3c). The D161^ECL2^-H165^ECL2^ interaction maintains its proton-sensing role but with increased distance (Fig. 3c). The D16^1.31^-E170^ECL2^-H269^7.36^ network shows tighter association compared to the G_s_-coupled GPR4 at pH 7.4 (Fig. 3c). The GPR4-G_q_ complex also displays distinct water molecule networks (Fig. 3d-e), suggesting G protein-specific diversity in proton sensing mechanisms.

**Fig. 3.**
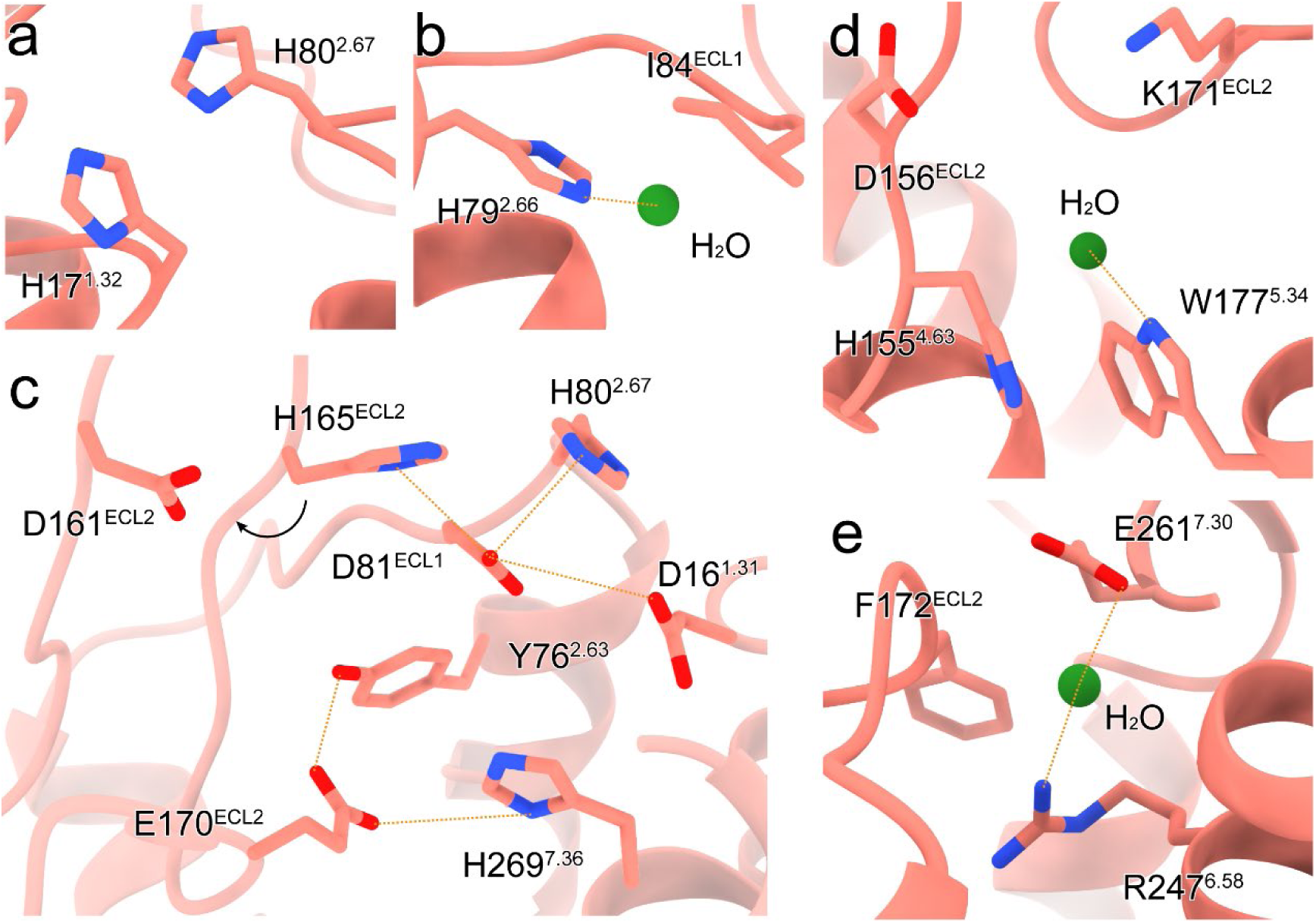
Proton recognition mode of GPR4 with G_q_ at physiobiological pH. **a** The preserved interactions between H17^1.32^ and H80^2.67^. **b** The interaction between H79^2.66^ and the common water molecule. **c** The extended polar network of H165^ECL2^ and H269^7.36^. **d-e** The special networks of the water molecules in G_q_-coupled GPR4. The water molecules are displayed as green spheres. The rotation direction of residues is labeled by a black arrow, and polar interactions are shown as orange dashed lines.

Beyond canonical proton-sensing residues, our analyses reveal crucial contributions from neutral phenylalanine and tyrosine residues (Fig. 4). The hydrophobic packing of V11^NT^, F167^ECL2^, F172^ECL2^, and F265^7.32^ facilitates ECL2 conformational shift toward N-terminus, enabling coordinated proton sensing by polar residues (Fig. 4a-c). The phenolic hydroxyl groups of Y76^2.63^ and Y98^3.33^ stabilize E170^ECL2^, a key component of proton-sensing networks (Fig. 2i, Fig. 3c, and Fig. 4d-f). Alanine mutation analysis of Y76^2.63^, Y98^3.33^, and F265^7.32^ supported by simulation results, confirms the functional significance of these interactions (Fig. 2j, Fig. 4g-h, and Fig. S9).

**Fig. 4.**
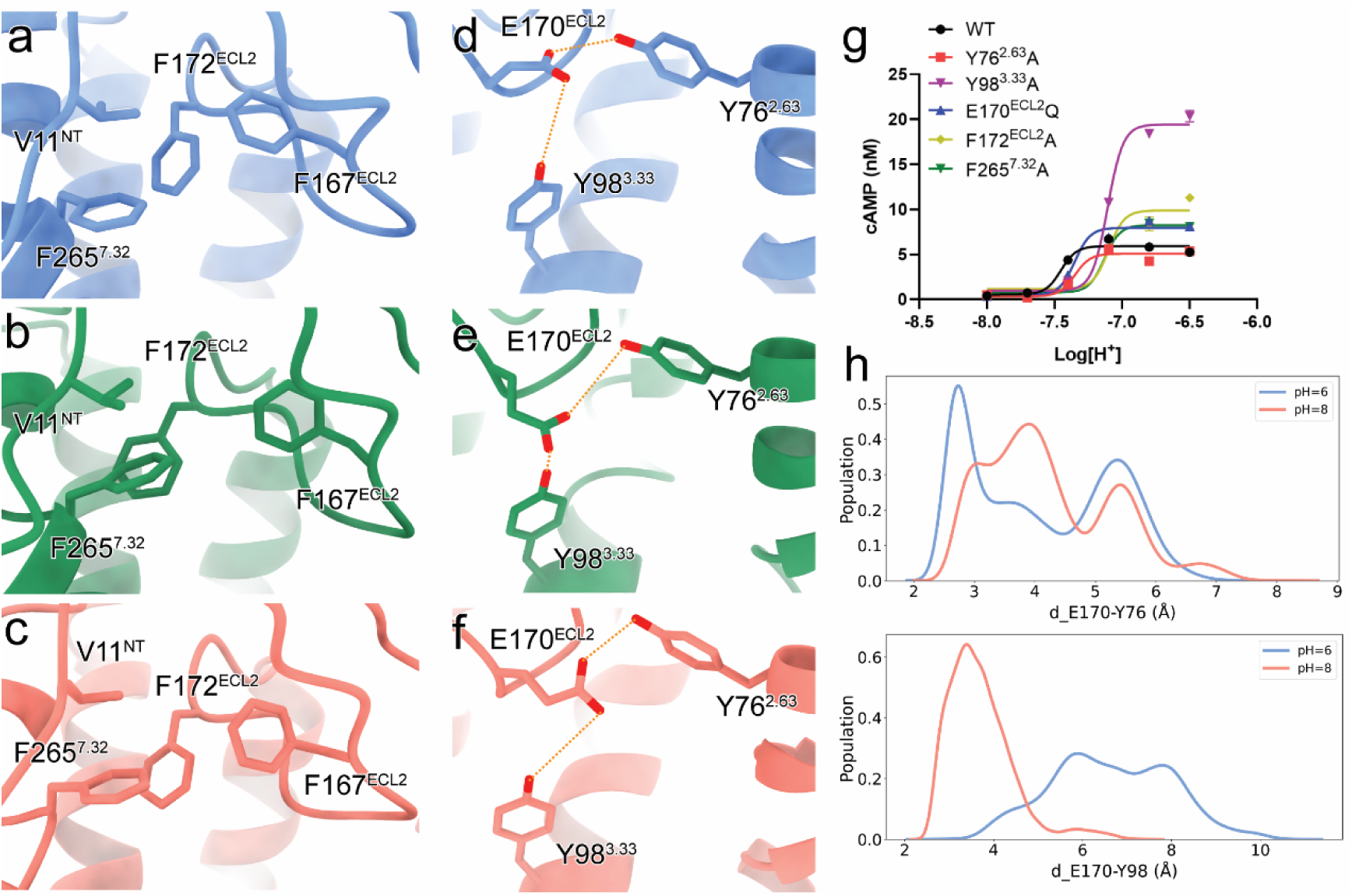
Involvement of neutral phenylalanine and tyrosine residues in proton sensing. a-c. Detailed interactions of phenylalanine residues in GPR4 complexes. **d-f** Detailed interactions of tyrosine residues in GPR4 complexes. Polar interaction are marked by orange dashed lines and colors are shown as indicated. **g** Effects of mutations on GPR4-induced cAMP accumulation at pH 6.5 (black bars) or 7.4 (red bars). The original data are provided in Supplementary Fig. S9. Values are shown as mean ± S.E.M. from three independent experiments. *P<0.05, **P<0.001 and ***P<0.0001 by one-way ANOVA followed by multiple comparisons test, compared with WT. **h** The conformation distribution under pH 6.0 and pH 8.0. The sidechain minimal distance distribution of Y76^2.63^-E170^ECL2^ (upper) and Y98^3.33^-E170^ECL2^ (lower).

### Proton-induced activation mechanism of GPR4

To decipher the molecular basis of pH-dependent GPR4 activation, we performed comprehensive structural comparisons among the fully active GPR4-G_s_ complex at pH 6.5, inactive β_2_AR structure (PDB: 2RH1), AlphaFold2-predicted GPR4 model, and simulated GPR4 at pH 8.0 (Fig. 5a).

**Fig. 5.**
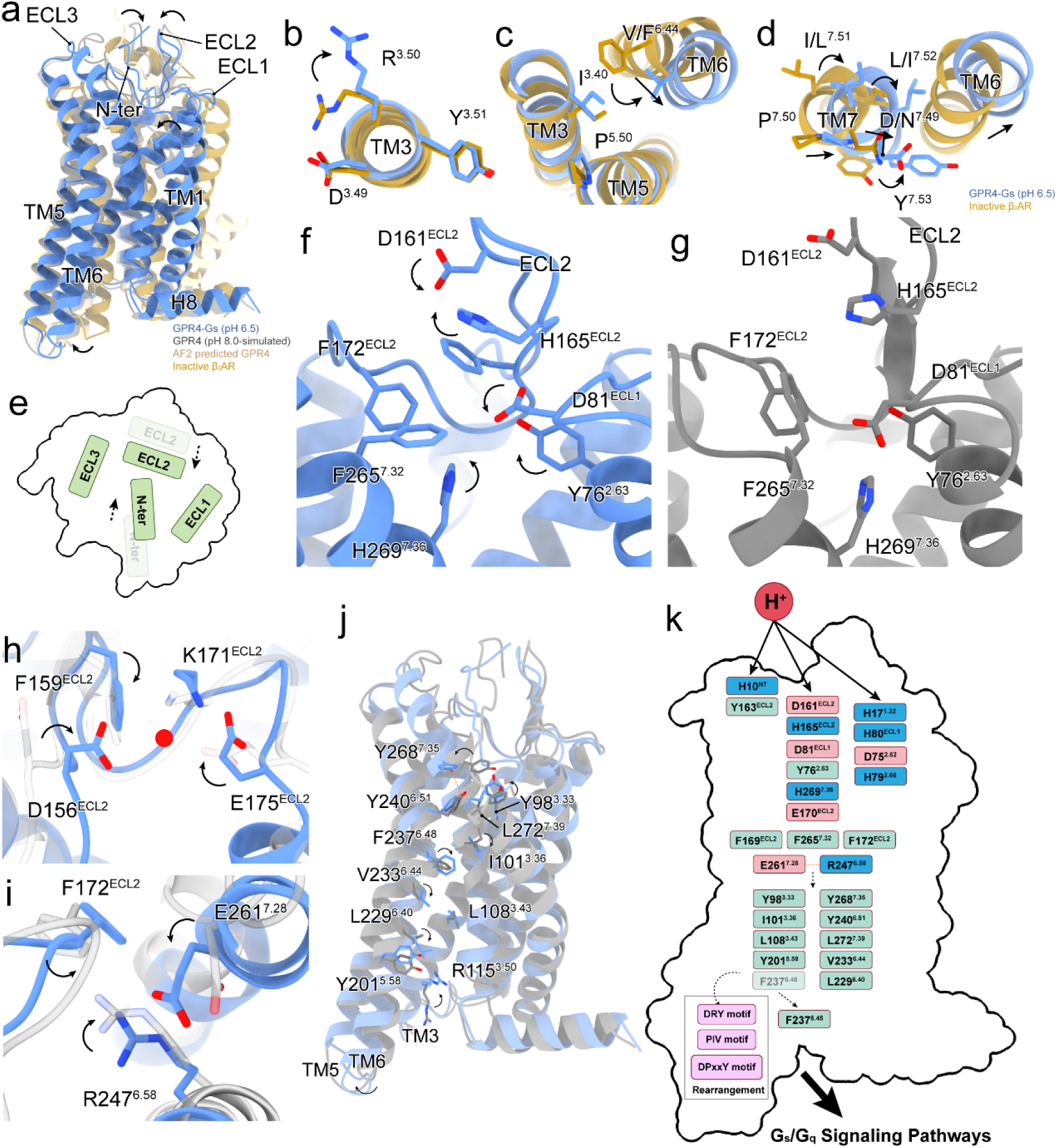
Proton-induced activation state of GPR4. **a** Overall structural comparison of GPR4 complexes, AlphaFold2 (AF2)-predicted GPR4 structure, the inactive structure of β_2_AR (PDB: 2RH1), and simulated GPR4 at pH 8.0. **b-d** The conformational changes of classical motifs in GPR4, including DRY (b), PIV (c), and DPxxY (d) motifs, compared with inactive β_2_AR. **e** Conformational changes of ECD upon activation. N-terminus and ECLs assemble to transduce activation signals. The transparent ones refer to their original positions and the normal ones refer to present positions. The black dashed arrows indicate displacement directions. **f-g** Detailed conformational changes of residues related to activation of GPR4-G_s_ at pH 6.5 and the simulated GPR4 at pH 8.0, the former is colored in blue and the latter is colored in gray. **h-i** Structural comparisons between the active and inactive (simulated) GPR4. **j** The potential propagation path for signal transduction. Related residues are highlighted. Black arrows represent the movements of the receptor and specific residues. **k** Schematic showing the proton-induced activation mechanism of GPR4. Direct interactions induced by protons are displayed by black arrows, and black dashed arrows indicate the connection of residues and conformational rearrangements of residues.

GPR4 exhibits characteristic GPCR activation features through rearrangements of conserved motifs (D^3.49^R^3.50^Y^3.51^/P^5.50^I^3.40^V^6.44^/C^6.47^F^6.48^xP^6.50^/D^7.49^P^7.50^xxY^7.53^) (Fig. 5b-d). Notably, GPR4 contains a unique DPxxY motif instead of the canonical NPxxY motif found in most class A GPCRs, a feature shared with other pH-sensing receptors GPR65 and GPR68 (Fig. S7). This distinctive motif appears crucial for pH responsiveness, consistent with previous findings that protonation states of D^2.50^ and D^7.49^ critically influence GPR4 activation ^26^.

Our structural analysis reveals that GPR4 activation initiates through significant conformational changes in ECD. N-terminus and ECLs undergo distinctive displacement and rotation, with activation signals propagating from N-terminus through ECL2 (Fig. 5a, e). The C9^NT^-C258^7.25^ disulfide bond serves as a critical anchor for N-terminus, establishing a fixed reference point for proton detection and ECLs recruitment (Fig. 2a). Mutations of these cysteines or N-terminus deletion severely compromise GPR4 function, highlighting their essential role (Fig. S6).

At both pH 6.5 and pH 7.4, key histidine residues (H10^NT^, H79^2.66^, H165^ECL2^, and H269^7.36^) in ECD undergo differential protonation in response to environmental pH. This protonation acts as a molecular switch for GPR4 activation by remodeling the electrostatic landscape of ECD and triggering receptor rearrangements (Fig. 5a, e). The polar network formed by D81^ECL1^, D161^ECL2^, and H165^ECL2^ induces repacking of a hydrophobic cluster (Y76^2.63^, F167^ECL2^, F172^ECL2^, and F265^7.32^), facilitating activation signal transduction (Fig. 5f-g). Subsequently, F159^ECL2^ repositions toward a water molecule that coordinates with D156^ECL2^, K171^ECL2^ and E175^ECL2^, stabilizing ECL2 conformation (Fig. 5h). R247^6.58^ rotates toward F172^ECL2^, establishing a salt-bridge with E261^7.28^(Fig. 5i). Core tyrosine residues, including Y98^3.33^ and Y268^7.35^, rotate toward Y240^6.51^ to propagate the signal downstream, triggering downward movements of I101^3.38^, L229^6.40^, and V233^6.44^ (Fig. 5j). Simultaneously, F237^6.48^ swings against TM3, facilitating the outward movement of TM6 (Fig. 5j). These conformational changes break the ionic lock through R115^3.50^-Y201^5.58^ interaction (Fig. 5j). Combined rearrangements of DRY, PIV, and DPxxY motifs open the intracellular pocket for G protein recruitment (Fig. 5j-k). Both simulation and mutagenesis studies validate the functional significance of these residues (Fig. S9, 11).

Our analyses reveal a sophisticated, multi-step activation mechanism (Fig. 5k). Protonation of key histidines initiates activation through ECD conformational changes, propagating through three networks involving H10^NT^, H79^2.66^, H165^ECL2^, and H269^7.36^ (Fig. 5k). While GPR4-Gα_s_ interfaces show minimal pH-dependent differences (Fig. S12a-e), the G_q_-coupled GPR4 at pH 7.4 exhibits slight differences, including αN rotation in Gα_q_ and α5 helix displacement (Fig. S12a-c). The GPR4-G_q_ structure shows unique E51^2.38^-D114^3.49^-Y358 polar network and modified ICL2-Gα interactions (Fig. S12d-f). This suggests that proton-induced activation of GPR4 is largely driven by external conformational shifts, rather than extensive alterations in the intracellular domain. This intricate mechanism provides a comprehensive molecular framework for understanding proton-induced activation of GPR4.

### Potential lipid regulation mechanism of GPR4

Our structural analysis revealed unexpected insights into lipid-mediated GPR4 regulation, particularly by LPC. While previous studies suggested the role of LPC role in GPR4 bioactivity ^57^, we identified distinct electron densities near classical allosteric sites adjacent to TM3-TM5 (Fig. 6a-c). These densities match the structural features of LPC, including its choline group, phosphate group, and short polar tail (Fig. 6a-c and Fig. S4). This observation aligns with previous studies ^35,36,57^ and our functional data showing ∼30% enhancement of GPR4 activation by LPC (Fig. 1c, 6e). A conserved water molecule positioned above the putative LPC binding site between TM4 and TM5 further defines this regulatory site (Fig. 6a-c).

**Fig. 6.**
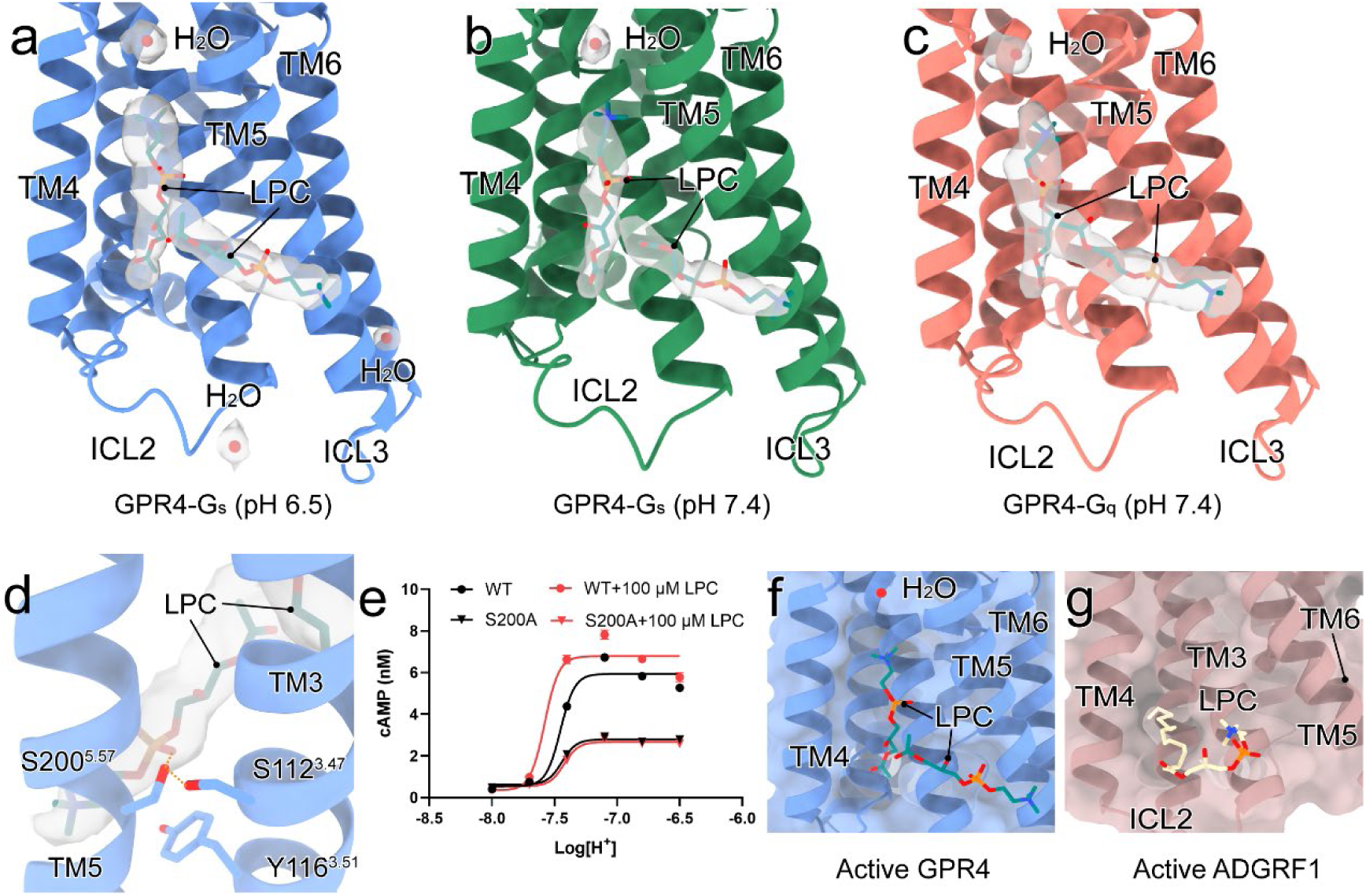
Novel lipid regulation of GPR4. a-c. The overall view of LPC-binding sites around GPR4-G_s_ at pH 6.5 (**a**)/7.4 (**b**), and GPR4-G_q_ at pH 7.4 (**c**), respectively. LPCs and water molecules are shown as sticks and red balls, respectively. **d** The engaged residues between LPCs and TM3/5 of GPR4. The polar interactions are displayed by orange dashed lines. **e** Measurement of S200A mutation effects on LPC positive allosteric activity (Analysis of cAMP accumulation pH-induced on GPR4 or mutants with or without 100 μM LPC). Values are represented as mean ± SEM of three independent experiments (n = 3). **f-g** Structural superposition of GPR4 and ADHRF1 (PDB: 7WU3).

The LPC binding pocket reveals key functional interactions. S200^5.57^ forms hydrogen bonds with both the phosphate group of LPC and S112^3.47^ (Fig. 6d). S200A^5.57^ mutation significantly reduces GPR4 activation and eliminates LPC responsiveness, confirming its crucial role (Fig. 6e). Y116^3.51^ of the conserved DRY motif also interacts with LPC, suggesting a potential mechanism for allosteric modulation (Fig. 6d). Computational analyses support the role of LPC, showing it has the lowest binding free energy among tested lipids (Fig. S13). This positive allosteric modulation by LPC parallels its recently reported effects on ADGRF1 ^51^. However, the partially overlapping binding sites between GPR4 and ADGRF1 suggest receptor-specific regulatory mechanisms (Fig. 6f-g).

## Discussion

The cell membrane functions as an environmental sensor, with proton concentration changes indicating various pathological conditions including cancer, metabolic disorders, and inflammation. Within the membrane pH-sensing components, proton-sensing GPCRs, including GPR4, contribute to cellular responses. Our study provides structural and mechanistic insights into proton-sensing mechanisms of GPR4, showing relationships between receptor structure, histidine protonation, water-mediated networks, and lipid regulation.

Recent studies have highlighted the critical role of ECD’s tight association and extended polar networks formed by histidine and acidic residues ^33,34^. The protonation of H^ECL2-45.47^ and H^7.36^ in non-human GPR4 activation ^33^ aligns well with our findings. Notably, H10^NT^, which emerged uniquely in mammalian ^33^, plays a central role in proton recognition. Through high-resolution cryo-EM structures, functional analyses, and computational simulations, we unveil a novel proton-sensing mechanism in human GPR4. Compared with the reported proton recognition of GPR68, GPR4 employs more extensive ECD residue networks, with key histidine protonation events serving as primary triggers for pH sensing and conformational stabilization. The conserved disulfide bond (C9^NT^-C258^7.25^) acts as a molecular anchor in both GPR4 and GPR68, facilitating the characteristic inward bending of N-terminus and TM1 critical for receptor function ^34^.

Our study identifies four key proton-sensing histidines - H10^NT^, H79^2.66^, H165^ECL2^, and H269^7.36^-whose differential protonation between pH 7.4 and 6.5 initiates precise conformational cascades. Similar to GPR68, GPR4 utilizes Y98^3.33^, E170^ECL2-45.52^, and H269^7.36^ to create a local polar environment facilitating signal transduction ^34^. We demonstrate that the proton-induced activation mechanism of GPR4 operates through three distinct networks centered on these key histidines in ECD. These triggers coordinated tyrosine rotations in the receptor core, propagating the activation signal through transmembrane domains to enable G protein recruitment. This intricate mechanism provides a comprehensive framework for understanding proton-induced GPCR activation.

Our structural analysis also unexpectedly revealed sophisticated lipid-mediated regulation of GPR4, particularly by LPC. Functional and computational analyses identify LPC as a positive allosteric modulator, suggesting an intricate interplay between lipid binding and histidine protonation that warrants further investigation. These findings have profound implications for both basic research and therapeutic development. The high expression and involvement in inflammation, angiogenesis, respiratory disorders, renal dysfunction, and cancer of GPR4 ^10,44,46^ position it as a promising therapeutic target. Our structural insights provide a foundation for structure-based drug design, as exemplified by existing antagonists like NE52-QQ57 and antagonist 3b ^43–46^. Future drug development could exploit the unique histidine-mediated pH-sensing mechanism, potentially through compounds that modulate histidine protonation states or their downstream effects.

In summary, our study establishes a molecular framework for the proton sensing and activation mechanisms of GPR4, showing interactions between water-mediated networks, histidine protonation, and lipid regulation. These findings may contribute to understanding pH-related pathologies and developing targeted therapeutics. Combining structural information with computational approaches could support the development of GPR4-targeted compounds for conditions involving pH dysregulation.

## Acknowledgments

The Cryo-EM data of GPR4-G_s_/G_q_ complexes were collected at the Advanced Center for Electron Microscopy, Shanghai Institute of Materia Medica (SIMM). The authors thank the staff at the Advanced Center for Electron Microscopy for their technical support. This work was partially supported by the Ministry of Science and Technology (China) grants 2018YFA0507002 (H.E.X.); the National Natural Science Foundation of China (82121005 to X.X., Y.J., and H.E.X., 32130022 to H.E.X., 82330113 to X.X., 82304579 to S.G., and 32171187 to Y.J.); the Shanghai Municipal Science and Technology Commission Major Project 2019SHZDZX02 (H.E.X.); the CAS Strategic Priority Research Program XDB37030103 (H.E.X.); the Lingang Laboratory (LG-GG-202204-01 to Y.J. and H.E.X.); the Shanghai Post-doctoral Excellence Program (2022684 to S.G.).

## Author Contributions

C.Y. and T.Z. screened the expression constructs, optimized the GPR4-G_s_/G_q_ complexes, prepared the protein samples for final structure determination, C.Y. participated in cryo-EM grid inspection, data collection, and model building; C.Y., T.Z., Z.L., S.G., and W.X. optimized the method for functional assays. C.Y., S.G., and W.X. designed the mutations and S.G., and X.W. executed the functional studies; X.X. supervised all the cellular experiments; C.Y. and Y.X. build and refine the structure models; X.H. participated in the molecular docking analysis; T.G, Z.L., Z.Z., Y.Z., Y.L., and X.C. participate in the experiments; H.E.X. conceived the project, initiated collaborations with X.X., and Y.J. conceived and supervised the project; C.Y. prepared the figures and drafted manuscript; H.E.X., X.X., and Y.J. wrote the manuscript with inputs from all authors.

## Methods

### Cells

Human embryonic kidney (HEK) 293 cells were obtained from ATCC (Manassas, VA, USA). High Five (Hi5) cells were purchased from Invitrogen.

### Construct

The wild type (WT) human GPR4 construct was cloned into the pFastBac^TM^ 1 vector (Thermo Fisher Scientific) with the N-terminal haemagglutinin signal peptide (HA) followed by a Flag tag and a 10×His tag. To enhance surface expression of GPR4, cytochrome b_562_RIL (BRIL) ^58^ followed by a tobacco etch virus (TEV) protease site was inserted into the N-terminus of GPR4. In order to further strengthen the coupling stability of GPR4 and G protein subunits, NanoBiT strategy is applied. In details, LgBiT fragment was fused to the C-terminus of GPCR with an optimized glycine-serine (GS) linker (GSSGGGGSGGGGSSG). Engineered Gα_s_ and Gα_q_ were used to improve the stability of GPR4 complexes. Gα_is_ is modified based on human Gα_s_ by replacing αN (MGCLGNSKTEDQRNEEKAQREANKK) with corresponding sequences of Gα_i_ (MGCTLSAEDKAAVERSKM) to facilitate scFv16 binding. Gα_isq_ is modified based on the miniGα_s_ scaffold and the its N-terminus replaced by corresponding sequences of Gαi1 (MGCTLSAEDKAAVERSKM). Human Gβ1 fused with a C-terminal 15-amino acid polypeptide linker (GSSGGGGSGGGGSSG) followed by a HiBiT and Gγ2 were cloned into the pFastBac^TM^ 1 vector, respectively. For cell-based functional assays in HEK293 cells, the wild-type GPR4 gene was subcloned into the pcDNA3.0 vector with the addition of an N-terminal HA tag. All the mutations used for functional studies were generated by QuickChange PCR and were verified by DNA sequencing.

### Expression and purification of Nb35

Nanobody-35 (Nb35) was expressed in Escherichia coli BL21 cells, and the cultured cells were grown in TB medium with 100 μg ml^−1^ ampicillin, 2 mM MgCl_2_, 0.1% glucose at 37 °C for 2.5 h until an optical density of 0.7–1.2 at 600 nm was reached. Then the culture was induced with 1 mM IPTG at 37 °C for 4–5 h, and the cells were collected and frozen at −80 °C. Nb35 was purified by nickel affinity chromatography, column or followed by overnight dialysis against 20 mM HEPES, pH 7.4, 100 mM NaCl, 10% glycerol. The Nb35 protein was verified by SDS–polyacrylamide gel electrophoresis and stored at −80 °C

### Expression of GPR4-G protein complexes

Hi5 cells were infected at a cell-density of 3.0×10^6^ cells per milliliter; five separate baculoviruses (GPR4, engineered Gα_s_ or Gα_q_, Gβ1-HiBiT, Gγ2, and scFv16) were co-added at a rational ratio of 1:1:1:1:1 into the insect cells. After culturing for 48 hours at 27℃, the cells were harvested by centrifugation. Then cell pellets were collected and stored at -80℃.

### GPR4-G_s_ and GPR4-G_q_ complexes formation and purification

Referring to the pH-dependent activation property of GPR4, we prepare two distinct buffers to maintain a stable pH by changing types and concentrations of buffer salt. For GPR4-G_s_ complex in acidic condition, cell pellets were resuspended and lysed in buffer containing 20 mM 2-Morpholinoethanesulphonic acid (MES) pH 6.5, 100 mM NaCl, 5 mM CaCl_2_ and 5 mM MgCl_2_, supplemented with EDTA-free complete protease inhibitor cock (APExBIO) and apyrase (25 mU/mL, Sigma). Complex formation was initiated during resuspension session and the suspension was incubated for 1 hour at room temperature. Then the supernatant was removed by centrifugation at 65,000 g for 40 min and resuspended sediment. Subsequently, 0.5% (w/v) lauryl maltose neopentyl glycol (LMNG, Anatrace) supplemented with 0.1% (w/v) cholesteryl hemisuccinate (CHS) (Anatrace) was added to solubilized GPR4 complex and extract it from membrane for 3 hours at 4℃. Insoluble material was then removed by centrifugation at 65,000 g for 40 min. The solubilized GPR4-G_s_ complex was incubated with Talon affinity resin overnight with 10 mM imidazole (pH 6.5) avoiding impurity binding. The resin was collected and washed with 20 column volumes of 20 mM MES pH 6.5, 100 mM NaCl, and a concentration gradient (12/15/18 mM) of imidazole (pH 6.5) and detergents (LMNG, GDN, and CHS). The complex was eluted with buffer containing 20 mM MES pH 6.5, 100 mM NaCl, 250 mM imidazole, 0.01% (w/v) LMNG and 0.002% (w/v) CHS, 0.005% (w/v) GDN and 0.001% (w/v) CHS. Finally, the complex was concentrated using a 15mL 100kDa cut-off Amicon Ultra Centrifugal Filter (Millipore) and Nb35 was added at a mole ratio of 1.5:1 incubating with GPR4 complex. The complex sample was then loaded onto a size exclusion chromatography on a Superose 6 Increase 10/300GL column (GE Healthcare) equilibrated with the buffer containing 20 mM MES pH 6.5, 100 mM NaCl, 0.00075% (w/v) LMNG and 0.00015% (w/v) CHS, 0.00025% (w/v) GDN and 0.00005% (w/v) CHS. The peak fractions of GPR4-G_s_ complex were collected and concentrated to 11.6 mg/mL using a 500 μL 100kDa cut-off Amicon Ultra Centrifugal Filter (Millipore) for making the cryo-EM grid. The purification process of GPR4-G_s_ and GPR4-G_q_ complexes at pH 7.4 is similar except for the substitution from 20 mM MES to 20 mM HEPES. The final purified samples of GPR4-G_S_ and GPR4-G_q_ at pH 7.4 were concentrated to 12.0 mg/mL and 5.2 mg/mL, respectively.

### Cryo-EM grid preparation and data collection

The purified samples of GPR4-G_s_ and GPR4-G_q_ complexes were applied onto holey carbon grids (Au300, R1.2/1.3, Quantifoil), which were glow-discharged at 25 mA for 50 s using PELCO easiGlow. Excess samples were blotted for 2s with Ted Pella filter papers (catalog number: 47000-100) under 100% humidity at 4℃. Afterward, the grids were vitrified by plunging into liquid ethane using a Vitrobot Mark IV (Thermo Fisher Scientific). For GPR4-G_s_ at pH 7.4 complex, cryo-EM data collection was performed on a Titan Krios G4 equipped with a Gatan K3 direct electron detector at 300 kV with a magnification of 105,000, corresponding to a pixel size 0.824 Å. Image acquisition was performed with EPU Software (Thermo Fisher Scientific, Eindhoven, Netherlands). A total of 3,751 movies were obtained at a total dose of 50 e^-^ Å^-2^ over 2.5 s exposure. For GPR4-G_s_ at pH 6.5 and GPR4—G_q_ at pH 7.4 complexes, cryo-EM data collection was performed on a Titan Krios G4 equipped with a Gatan Quantum-LS Energy Filter (GIF) and a Falcon 4 direct electron detector at 300 kV with a magnification of 165,000, corresponding to a pixel size 0.73 Å. Image acquisition was performed with EPU Software. A total of 6,696 and 7,848 movies were obtained at a total dose of 50 e^-^ Å^-2^ over 3 s exposure, respectively. All movies were collected at the Advanced Center for Electron Microscopy at Shanghai Institute of Materia Medica, Chinese Academy of Sciences.

### Cryo-EM data processing

All dose-fractionated image stacks were subjected to beam-induced motion correction by RELION4.0 ^59^. The defocus parameters were estimated by CTFFIND4.1 ^60^. The following data processing was performed using RELION4.0 and CryoSPARC4.4.1, respectively.

For the GPR4-G_s_ complex at pH 6.5, data processing was performed in RELION4.0. Particle selection yielded 2,798,739 particles, which were subjected to reference-free 2D classification. The map of LY3154207-DRD1-G_s_ (PDB: 7CKZ) ^61^ low-pass-filtered to 40 Å was used as an initial reference model for 3D classification and 2,312,673 particles were selected to do further processing. Then, multiple rounds of 3D classifications produced one high-quality subset accounting for 596,409 particles. These particles were subsequently subjected to 3D refinement, post-processing, and deepEMhancer ^62^, which generated a map with an indicated global resolution of 2.36 Å at an FSC of 0.143.

For the GPR4-G_s_ complex at pH 7.4, data processing was performed in CryoSPARC4.4.1. Particle selection yielded 3,349,414 particles, which were subjected to reference-free 2D classification. After rounds of 2D classification, 206,260 bad-quality particles were subjected to ab-initio reconstruction and produced three distinct density maps. Then, we imported the map of solved GPR4-G_s_ complex (pH 6.5) and combined them as input 3D references for hetero refinement. One obvious high-quality subset of 626,376 particles was chosen to do further optimization, including global/local CTF refinement, homo/non-uniform refinement, local refinement, and deepEMhancer. A map with an indicated global resolution of 2.59 Å at an FSC of 0.143 was generated.

For the GPR4-G_q_ complex at pH 7.4, data processing was performed in CryoSPARC4.4.1. Particle selection yielded 4,555,181 particles, which were subjected to reference-free 2D classification. After rounds of 2D classification, 245,566 bad-quality particles were subjected to ab-initio reconstruction and produced three distinct density maps. Then, we imported the map of solved GPR4-G_s_ complex (pH 6.5) and combined them as input 3D references for hetero refinement. One obvious high-quality subset of 550,055 particles was chosen to do further optimization, including global/local CTF refinement, homo/non-uniform refinement, local refinement, and deepEMhancer. a map with an indicated global resolution of 2.55 Å at an FSC of 0.143 was generated.

### Methods of MD simulations

The simulation systems were derived from the GPR4-Gs protein complex at pH 6.5, with G proteins removed prior to simulations. Acetyl (ACE) and N-methyl (NME) groups were added using PyMOL as described in reference ^63^. The GPR4 was embedded into a 75x75 Å POPC lipid bilayer using packmol-memgen software ^64^, surrounded by a 15 Å aqueous layer. Ionic strength was adjusted to 0.15 mol/L NaCl, with additional counterions. We utilized the FF19SB, Lipid21, and GAFF2 force fields for amino acids, lipids, and ligands, respectively ^65–67^. Systems underwent a minimization and a six-step equilibration process following the CpHMD prep protocol (https://gitlab.com/shenlab-amber-cphmd/cphmd-prep). Three independent 500 ns production runs were conducted for each system using pmemd.cuda in Amber22 ^68^ under the NPT ensemble at 300 K and 1 atm. pH values of 6 and 8 were maintained in all-atom mode. Long-range electrostatic interactions were managed using the Particle Mesh Ewald method, while short-range electrostatic and van der Waals interactions were handled with a 12 Å cutoff, transitioning smoothly between 10 and 12 Å. SHAKE was applied to constrain the bonds containing hydrogens, permitting the timestep of 2 fs. Minimal distances were calculated using the “nativecontact” command in CPPTRAJ^69^.

### Binding free energy calculation

Molecular Mechanics/Generalized Born Surface Area (MM/GBSA) was applied for binding free energy estimation. For MM/GBSA calculations, analogues of LPC were generated based on the GPR4 complex at pH 6.5 using the builder module in PyMOL. These complexes were subsequently prepared using the Protein Preparation Wizard in Schrödinger’s Maestro. During this process, bond orders were assigned, hydrogens were added to the protein, disulfide bonds were created, and residue heteroatom states were defined using Epik at pH 6.5. Each complex was then minimized using the OPLS4 force field, applying a 3.0 Å constraint on heavy atoms. An implicit membrane was added according to the helix orientation, with a thickness of 44.5 Å. Finally, Prime MM-GBSA calculations were performed using the VSGB solvation model and the OPLS4 force field.

### pK_a_ calculation

The pK_a_ values of histidine residues were calculated using the PROPKA server available at https://www.ddl.unimi.it/vegaol/propka.html ^70^. The PROPKA algorithm estimates residue-specific pK_a_ values based on structural and environmental factors within protein models. Input structures of the protein were prepared in PDB format. The resulting pK_a_ values for histidines were extracted and plotted to compare the protonation behavior of histidine residues under these conditions. No additional modifications or parameter adjustments were made to the default settings of the PROPKA server.

### Cell culture and transfection

HEK293 cells were obtained from ATCC (Manassas, VA, USA) and cultured in DMEM supplemented with 10% (v/v) FBS, 100 mg/L penicillin, and 100 mg/L streptomycin in 5% CO_2_ at 37 ℃. For transient transfection, approximately 2.5×10^6^ cells were mixed with 1 µg plasmids in 200 µL transfection buffer, and electroporation was carried out with a Scientz-2C electroporation apparatus (Scientz Biotech, Ningbo, China). The experiments were carried out 24 hours after transfection.

### Stimulus buffer

Experiments were carried out in a physiological salt solution (PSS) containing 130 mM NaCl, 0.9 mM NaH_2_PO_4_, 5.4 mM KCl, 0.8 mM MgSO_4_, 1.0 mM CaCl_2_, 25 mM glucose. This solution was buffered with HEPES/EPPS/MES (8 mM each; HEM-PSS), to cover a wider pH range. The pH of all solutions was adjusted using a carefully calibrated pH meter (Mettler Toledo). All data in this report are referenced to pH at room temperature.

### cAMP accumulation assay

Intracellular cAMP levels were detected with an HTRF cAMP kit obtained from PerkinElmer, according to the manufacturer’s instructions. In brief, HEK293 cells transfected with GPR4 WT receptor or its mutations were seeded at a density of 4×10^4^ per well into 96-well culture plates and incubated for 24 h at 37 ℃ in 5% CO_2_. The next day, the cells were incubated for 30 min under the different pH level in the presence of IBMX at room temperature. The reaction was terminated by adding 100 µL lysis buffer and product was diluted in the appropriate proportion. Then plated 10 µL lysis product onto 384-well assay plates. Another 10 µL lysis buffer containing ULight-anti-cAMP and Eu-cAMP tracer was added. After 60 minutes incubation in the dark, HTRF signals were detected with an Envision 2101 plate reader (PerkinElmer, Waltham, MA). The cAMP concentrations were calculated from standard curves.

### LPC dose curve

HEK293 cells and transfected with GPR4 WT receptor were seeded at a density of 4×10^4^ per well into 96-well culture plates and incubated for 24 h at 37 ℃ in 5% CO_2_. The next day, cells were incubated with 50 µL of pH7.4 stimulus buffer containing LPC at various concentrations for 30 minutes in the presence of IBMX at room temperature. The reaction was terminated by adding 100 µL lysis buffer and product was diluted in the appropriate proportion. Another 10 µL lysis buffer containing ULight-anti-cAMP and Eu-cAMP tracer was added. After 60 minutes incubation in the dark, HTRF signals were detected with an Envision 2101 plate reader (PerkinElmer, Waltham, MA). The cAMP concentrations were calculated from standard curves.

---

**Fig. S1.**
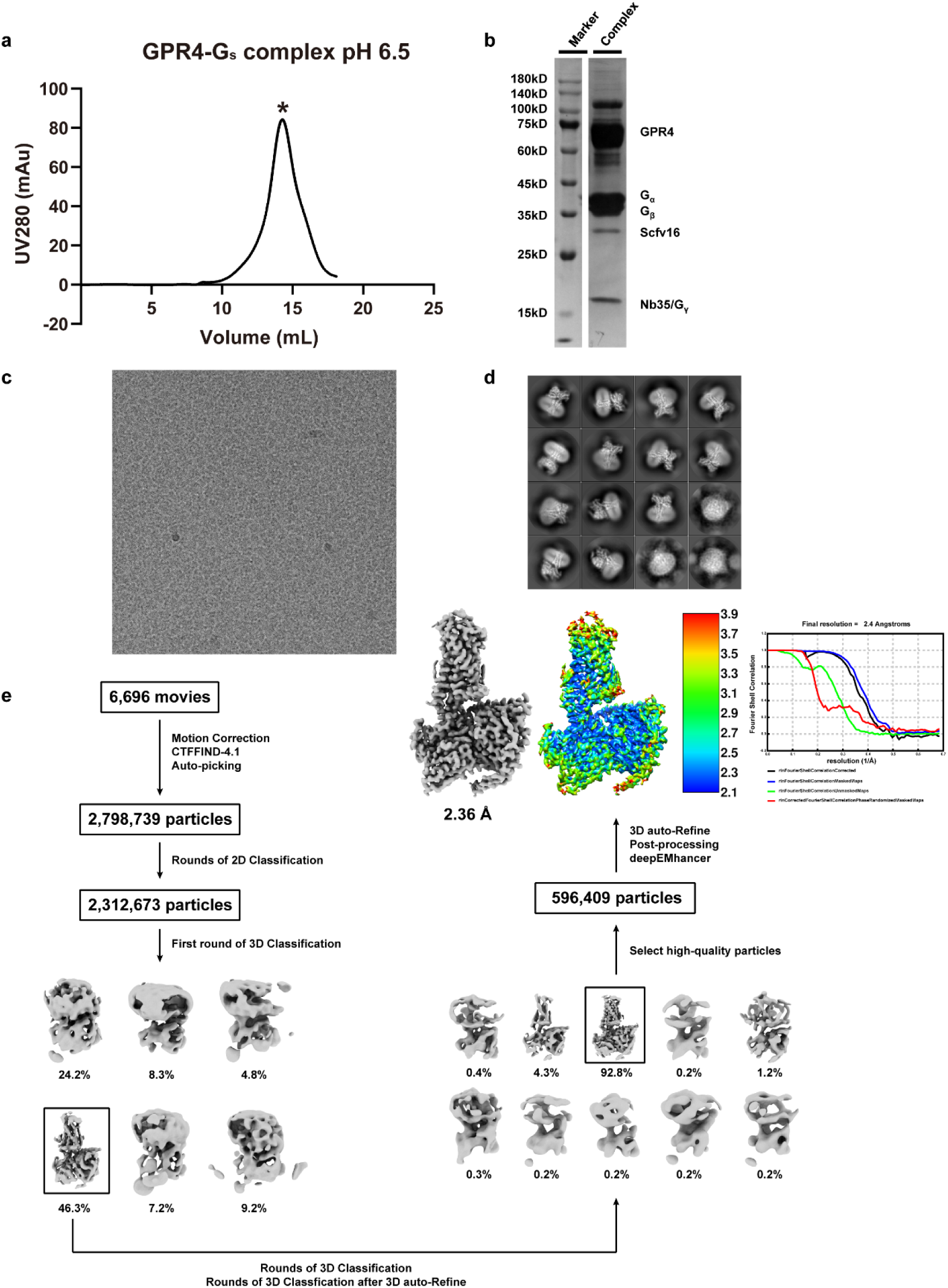
Cryo-EM data processing of GPR4-G_s_ at pH 6.5. a-b. Representative size-exclusion chromatography elution profile (**a**) and SDS-PAGE analysis (**b**) of GPR4-G_s_ at pH 6.5. **c-d** Representative cryo-EM micrograph (**c**) and representative 2D average classes are shown (**d**). Scale bar, 50 nm. **e** Flowchart of cryo-EM data processing, cryo-EM maps and “Gold-standard” FSC curves. Local resolution map is generated by ResMap.

**Fig. S2.**
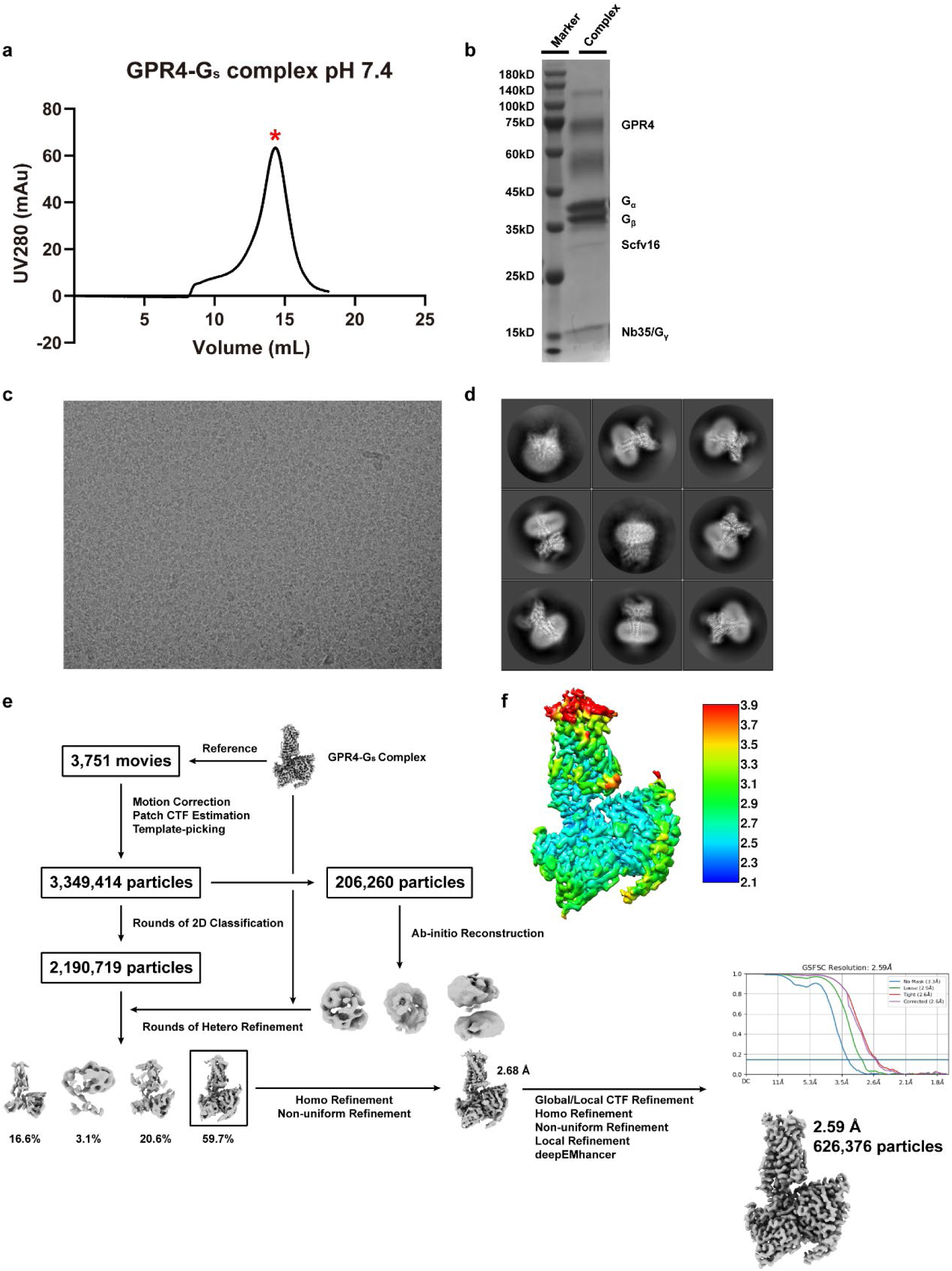
Cryo-EM data processing of GPR4-G_s_ at pH 7.4. a-b. Representative size-exclusion chromatography elution profile (**a**) and SDS-PAGE analysis (**b**) of GPR4-G_s_ at pH 7.4. **c-d** Representative cryo-EM micrograph (**c**) and representative 2D average classes are shown (**d**). Scale bar, 50 nm. **e** Flowchart of cryo-EM data processing, cryo-ResMap.

**Fig. S3.**
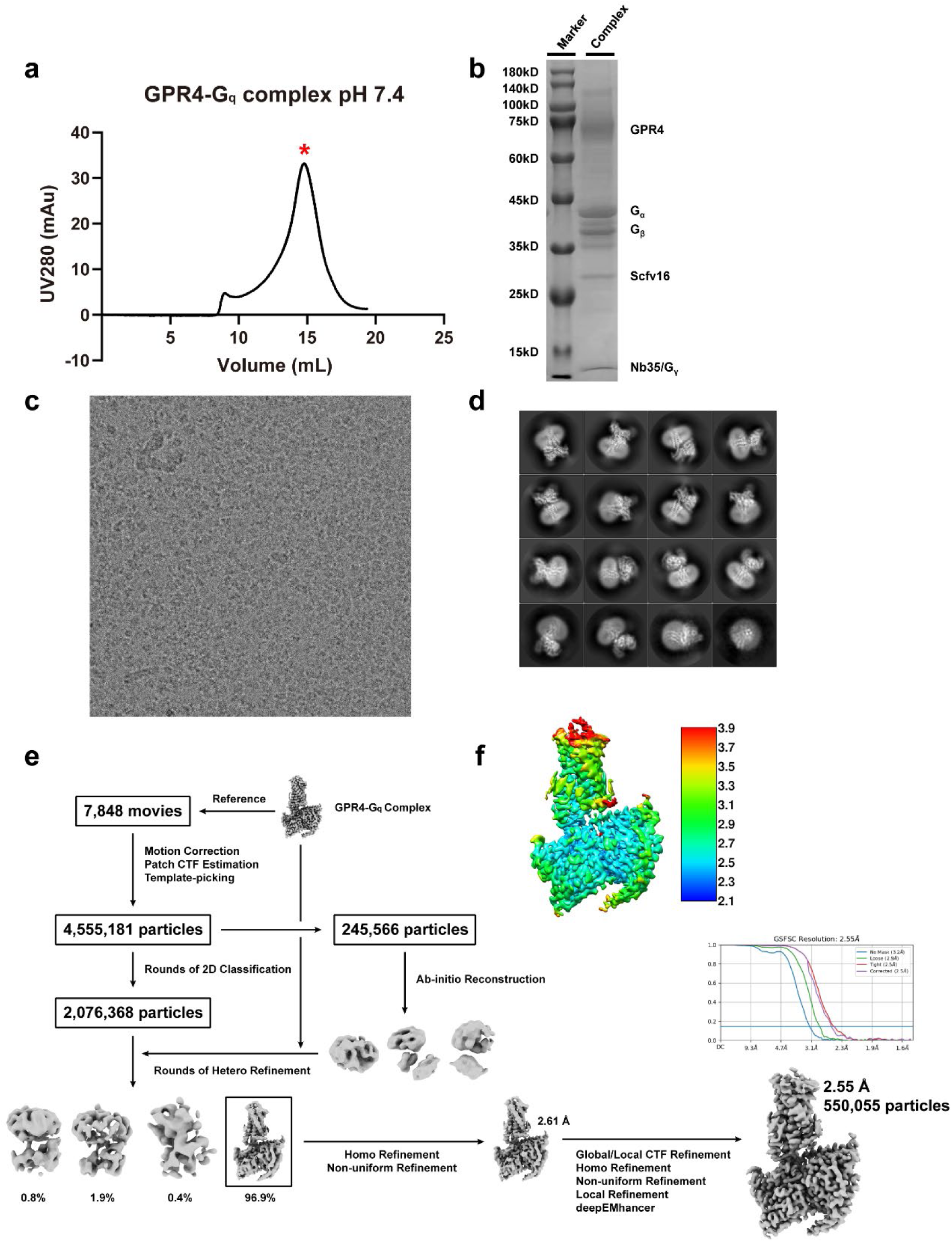
Cryo-EM data processing of GPR4-G_q_ at pH 7.4. a-b. Representative size-exclusion chromatography elution profile **(a**) and SDS-PAGE analysis (**b**) of GPR4-G_q_ at pH 7.4. **c-d** Representative cryo-EM micrograph (**c**) and representative 2D average classes are shown (**d**). Scale bar, 50 nm. **e** Flowchart of cryo-EM data processing, cryo-EM maps and “Gold-standard” FSC curves. Local resolution map is generated by ResMap.

**Fig. S4.**
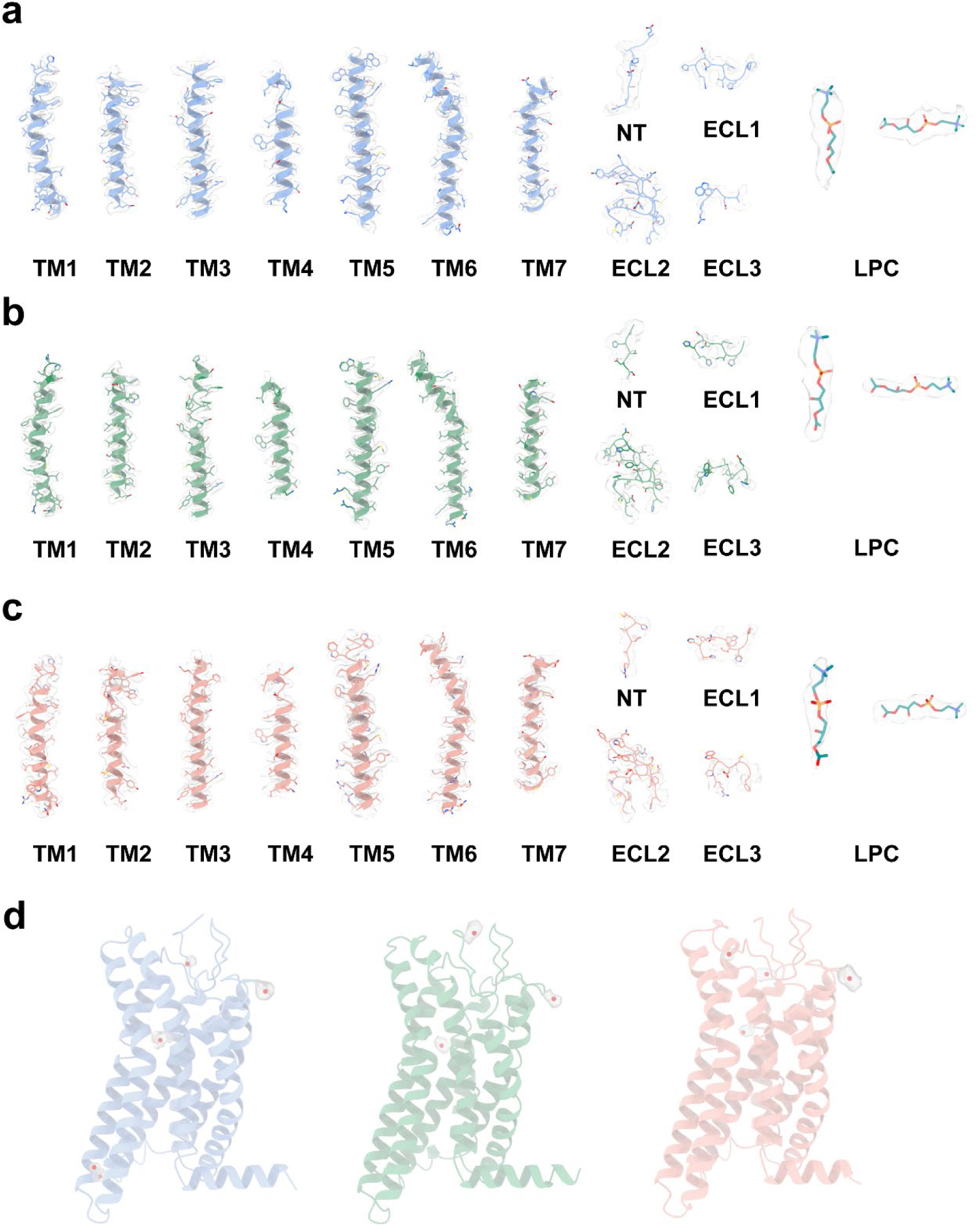
Representative cryo-EM density maps of GPR4 and lipids. Cryo-EM density maps of the seven transmembrane (TM) helices, N-terminus, ECLs, and bioactive lipids (LPC) for GPR4-G_s_ at pH 6.5 **(a)**, GPR4-G_s_ at pH 7.4 **(b)**, and GPR4-G_q_ at pH 7.4 **(c)**. **d** The densities of the water molecules of active GPR4 complexes. Colors are shown as indicated.

**Fig. S5.**
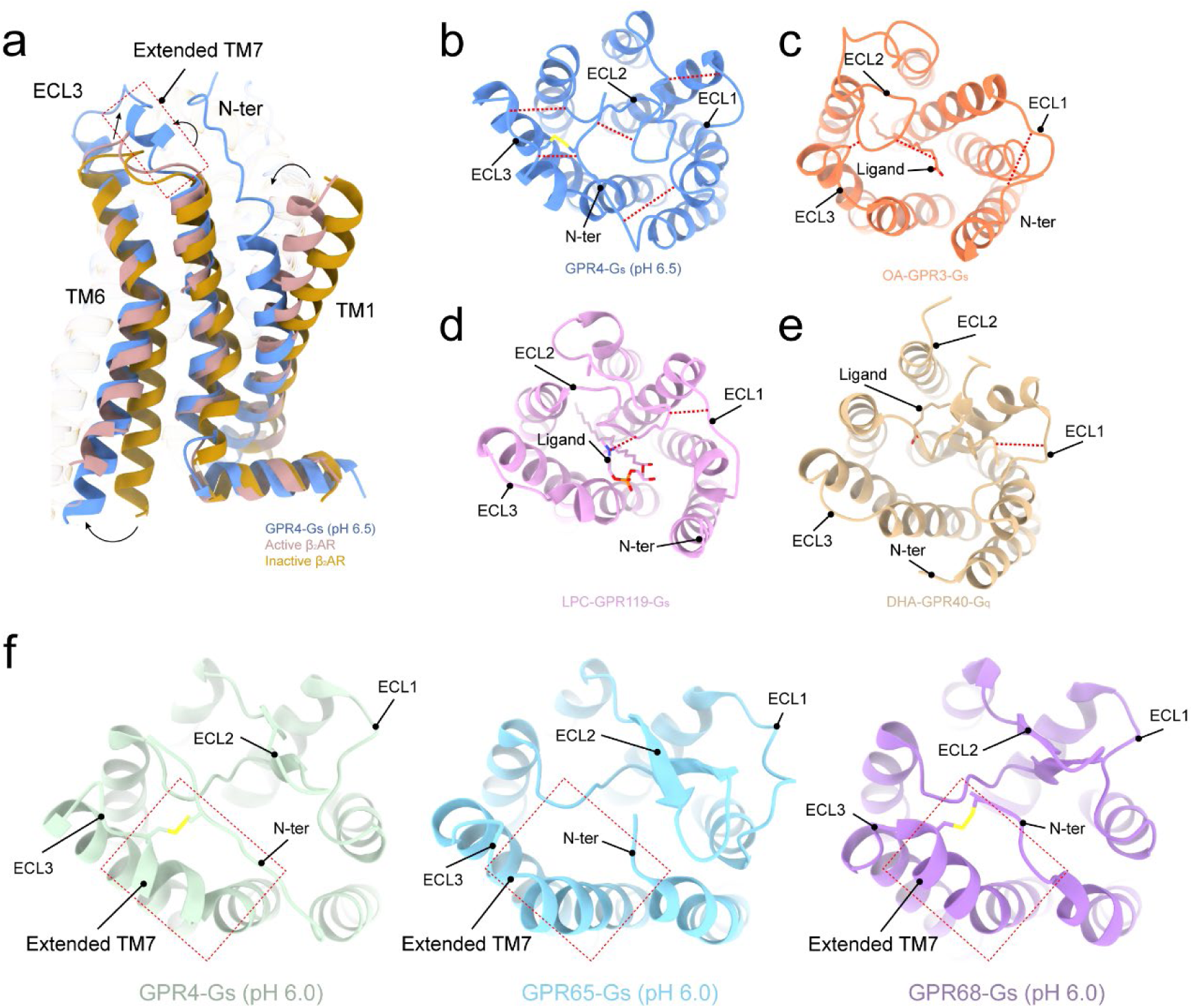
Structural comparison with representative GPCRs. **a** Structural comparison with active and inactive β_2_AR (PDB: 7DHI/2RH1), respectively. **b-e** Distinct ECD conformations of GPR4 and lipid-liganded GPCRs. The distances are shown in red dashed lines and directions of related displacement are displayed by black arrows. **f** The top views of published proton sensing GPCRs. The dashed red rectangles display the common structural features among GPR4, GPR65, and GPR68, including a special disulfide bond and an extended TM7. Colors are shown as indicated.

**Fig. S6.**
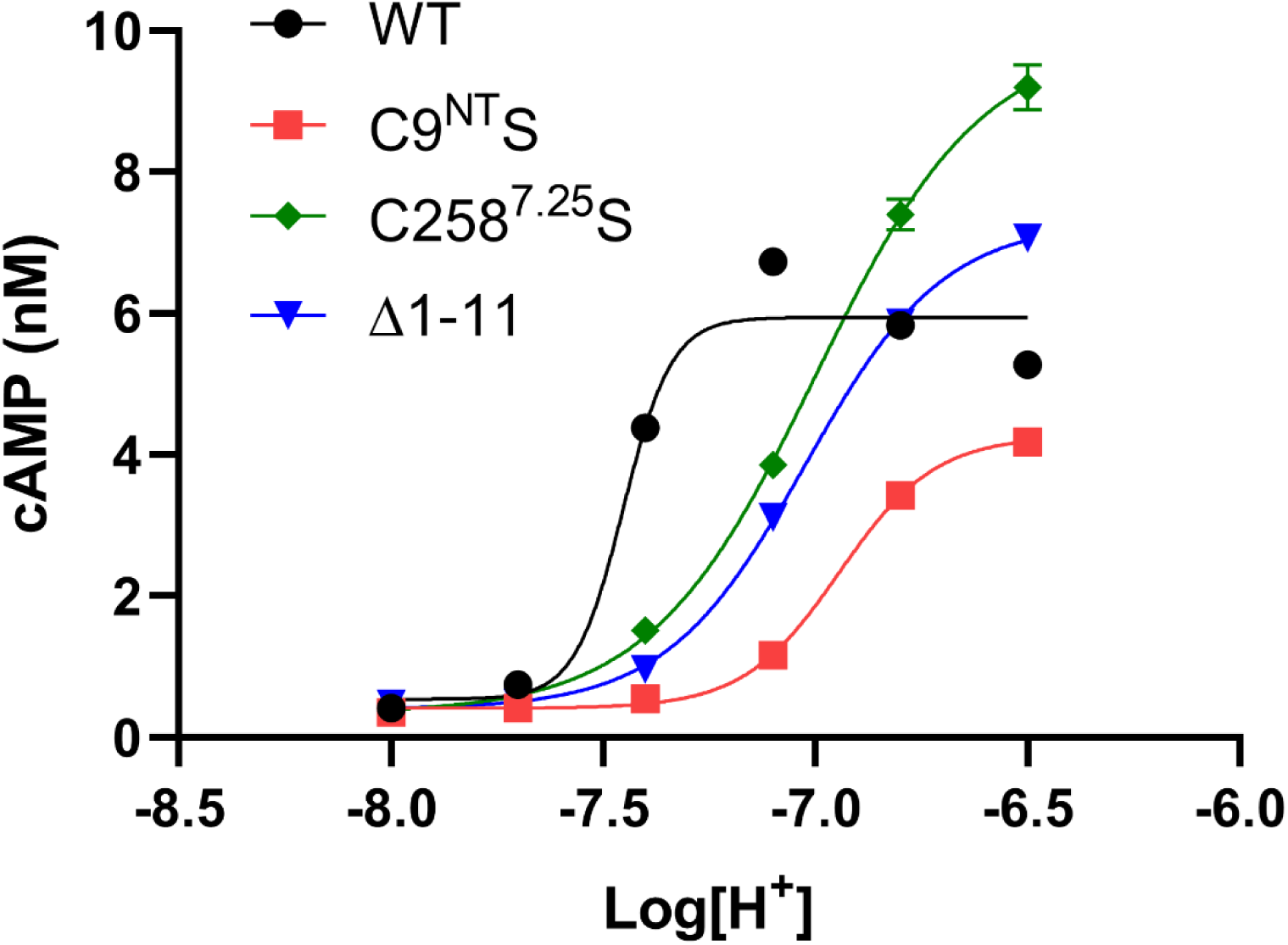
cAMP accumulation assay of pH-induced on GPR4 or mutants. Curves showing pH-dependent cAMP accumulation in cells overexpressing GPR4 or mutants. Values are represented as mean ± SEM of three independent experiments (n = 3).

**Fig. S7.**
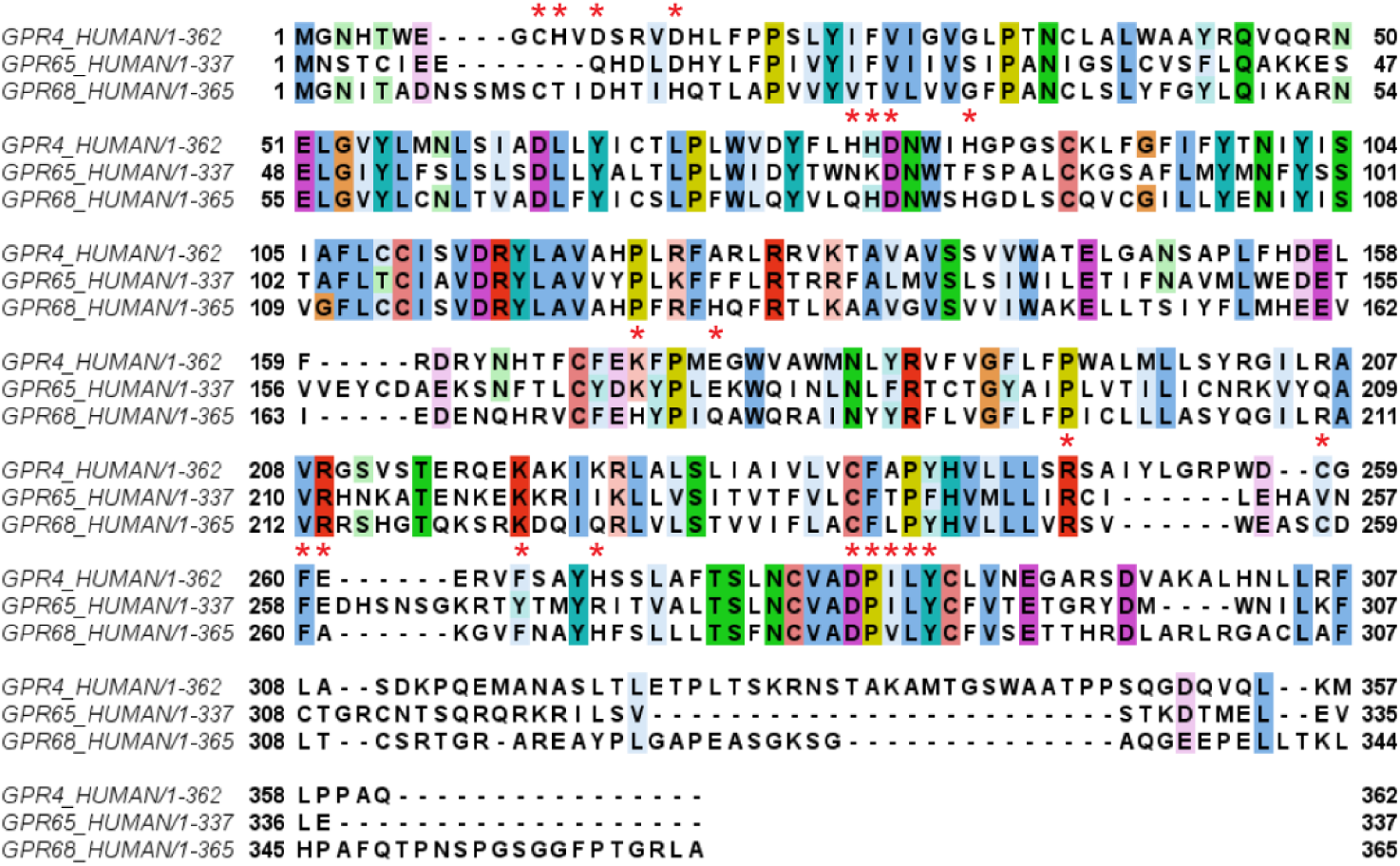
Sequence alignment of proton-sensing GPCR. Conserved sites are marked by red stars and sequence alignment is performed in Jalview by Mafft method.

**Fig. S8.**
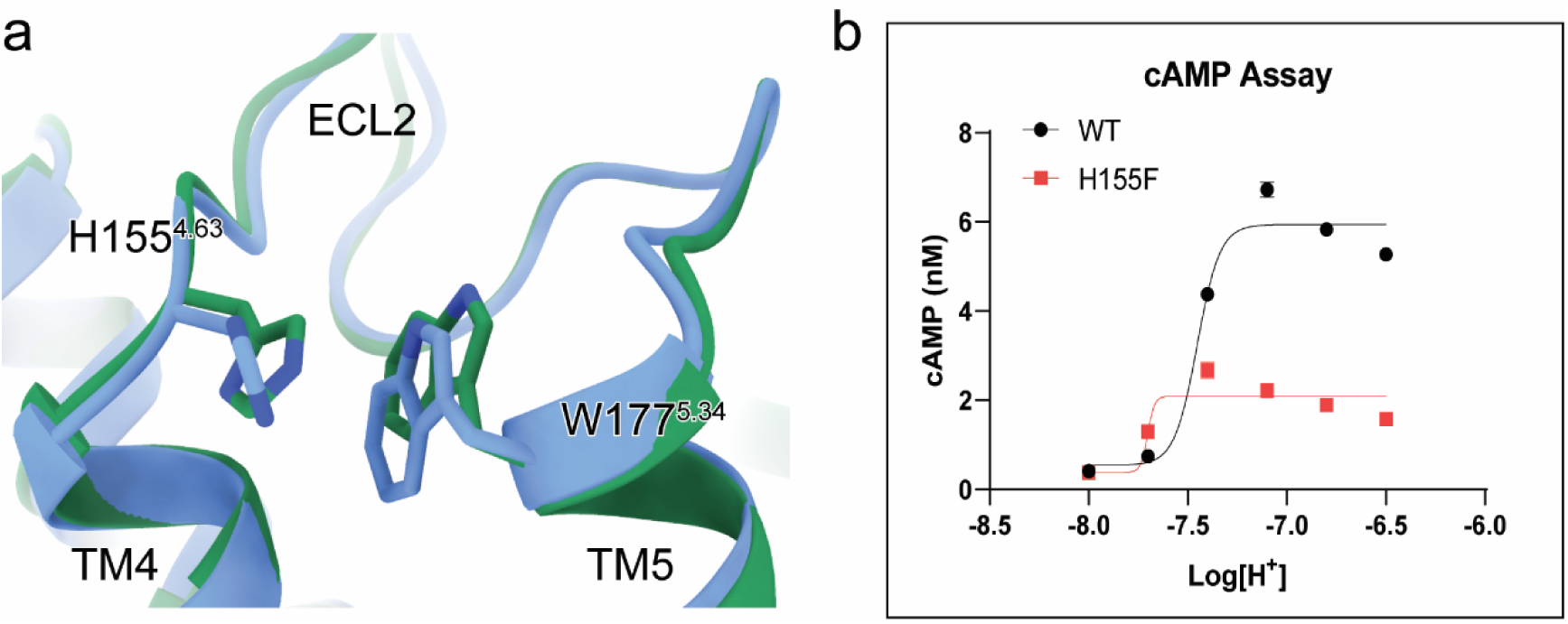
π-π stacking of H155^4.63^ and W177^5.34^. **a** The existence of the important π-π stacking between TM4 and TM5. **b** Effect of different pH on the WT and the H155F mutant of GPR4 using cAMP accumulation assay. Data shown are mean ± S.E.M. of three independent experiments.

**Fig. S9.**
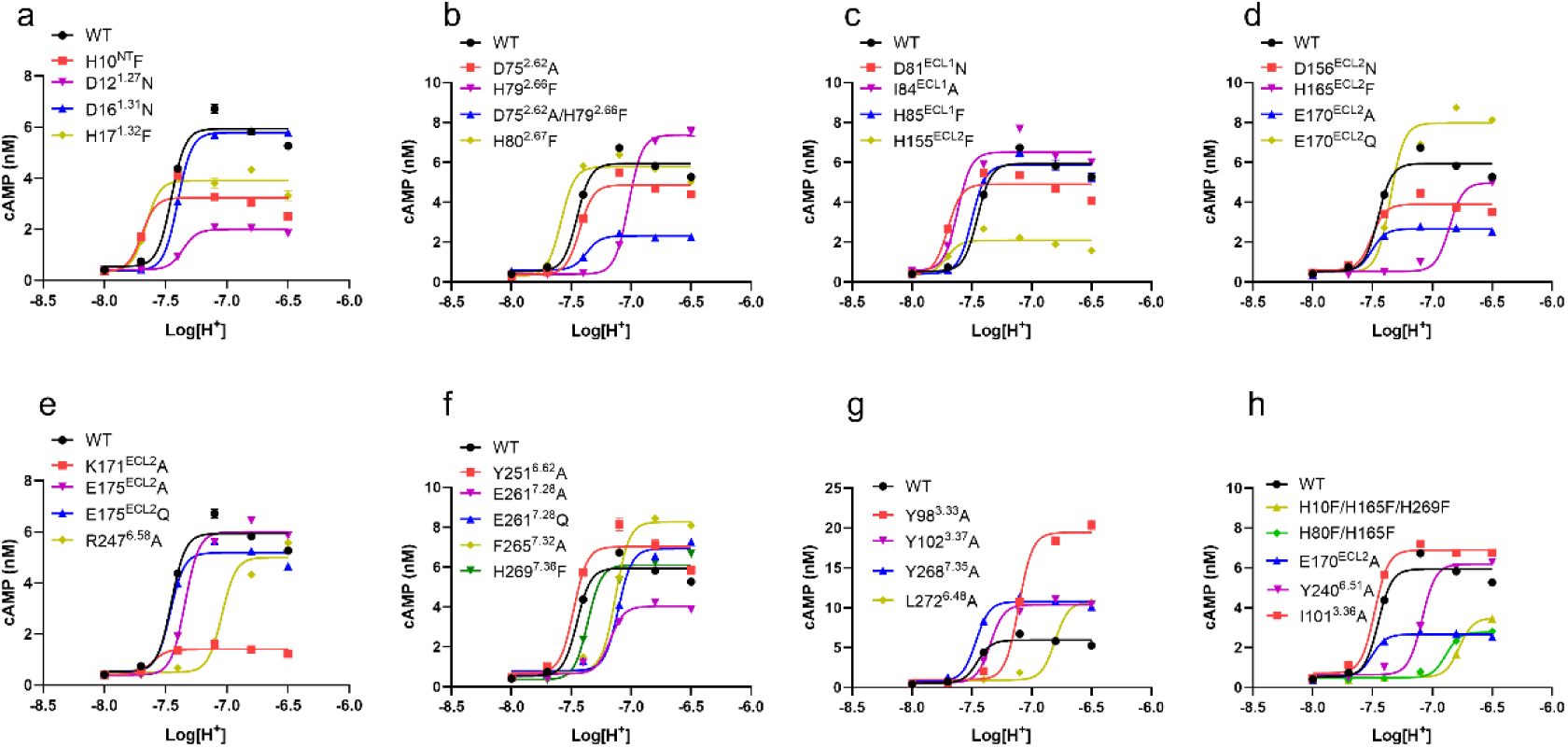
Cyclic AMP accumulation analysis for GPR4. Effects of different pH on the WT and mutants of GPR4 using cAMP accumulation assay. Data shown are mean ± S.E.M. of three independent experiments.

**Fig. S10.**
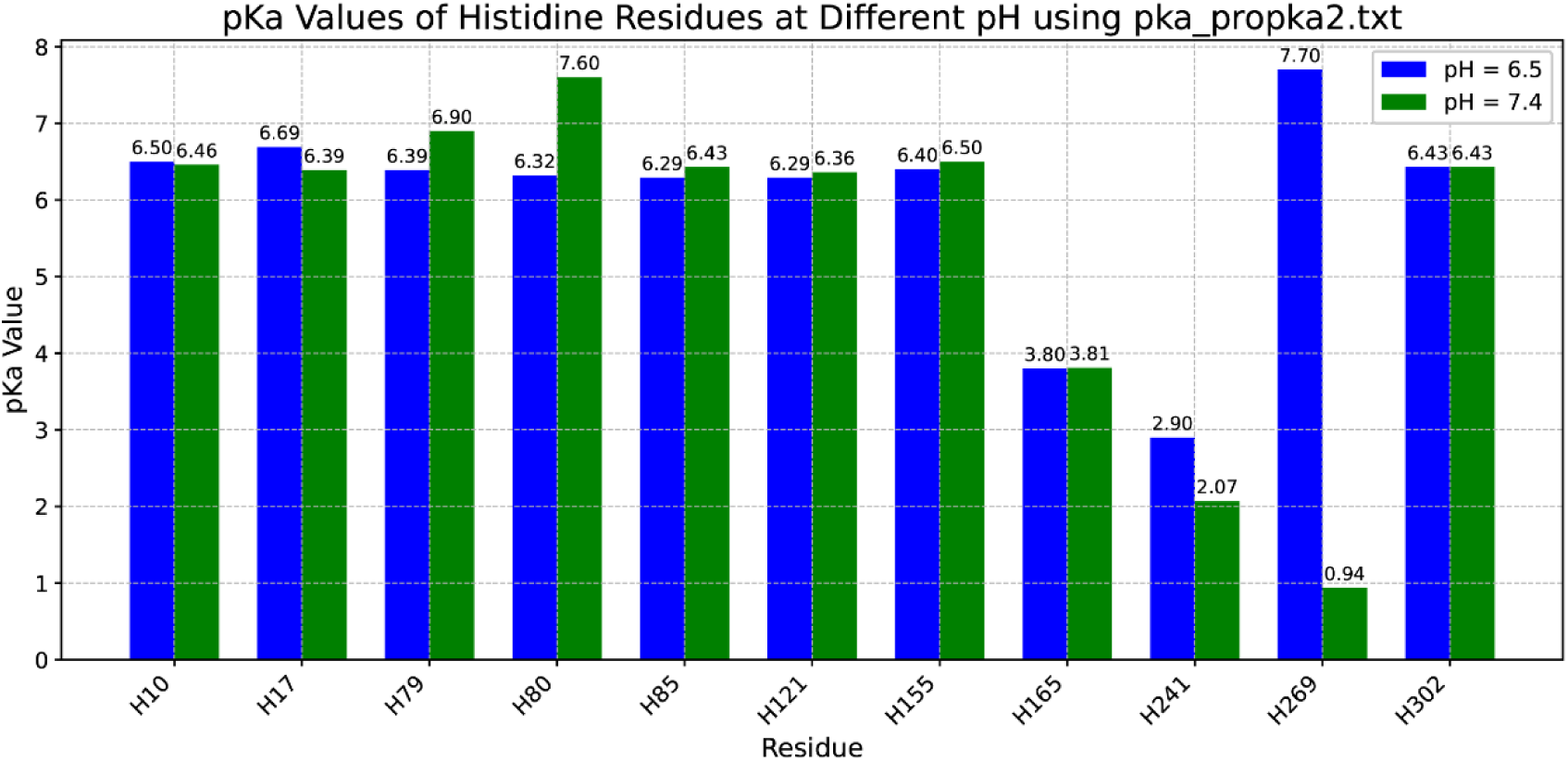
Calculations of pK_a_ for extracellular histidine residues by PROPKA. Residues are labeled on the x-axis, and their corresponding pK_a_ values are displayed above each bar.

**Fig. S11.**
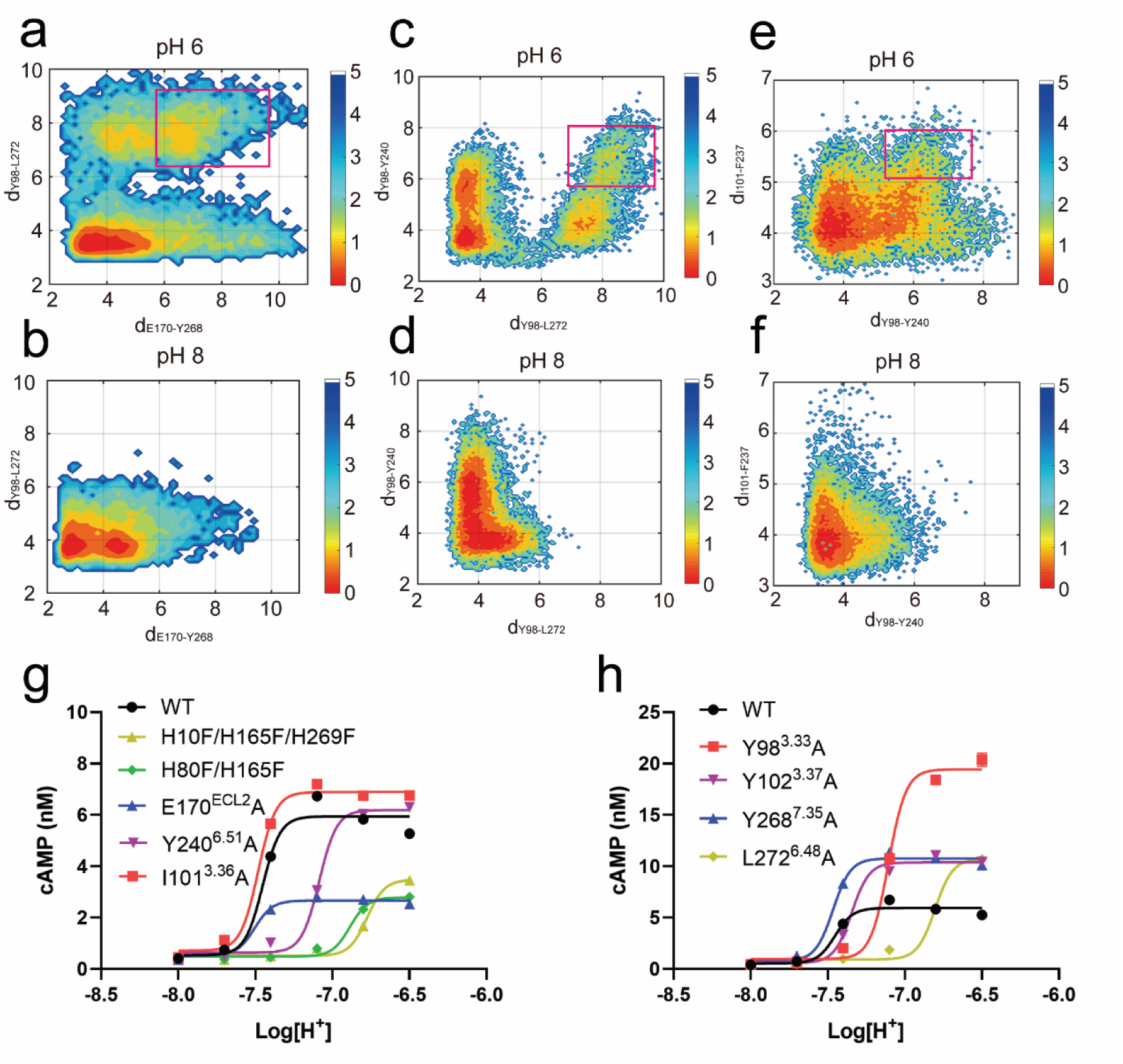
Evaluations of receptor activation mechanism exploration by molecular simulations and cAMP accumulation assay. conformation distribution under pH 6.0 and pH 8.0. a-f. MD simulations of GPR4 activation. **a-b** The free energy landscape of the minimal distance between sidechain of E170^ECL2^ and Y268^7.35^ (X axis) and the sidechain minimal distance between Y98^3.33^ and L272^7.39^ (Y axis) at pH 6.0 (a) and pH 8.0 (b), respectively. **c-d** The free energy landscape of the sidechain minimal distance between Y98^3.33^ and L272^7.39^ (X axis) and the sidechain minimal distance between Y983.33 and Y2406.51 (Y axis) at pH 6.0 (c) and pH 8.0 (d), respectively. **e-f** The free energy landscape of the minimal distance between Y98^3.33^ and Y240^6.51^ (X axis) and the sidechain minimal distance between I101^3.36^ and F237^6.48^ (Y axis) at pH 6.0 (e) and pH 8.0 (f), respectively. **g-h** Effects of different pH on the GPR4 (WT) and mutants using cAMP accumulation assay. Data shown are mean ± S.E.M. of three independent experiments.

**Fig. S12.**
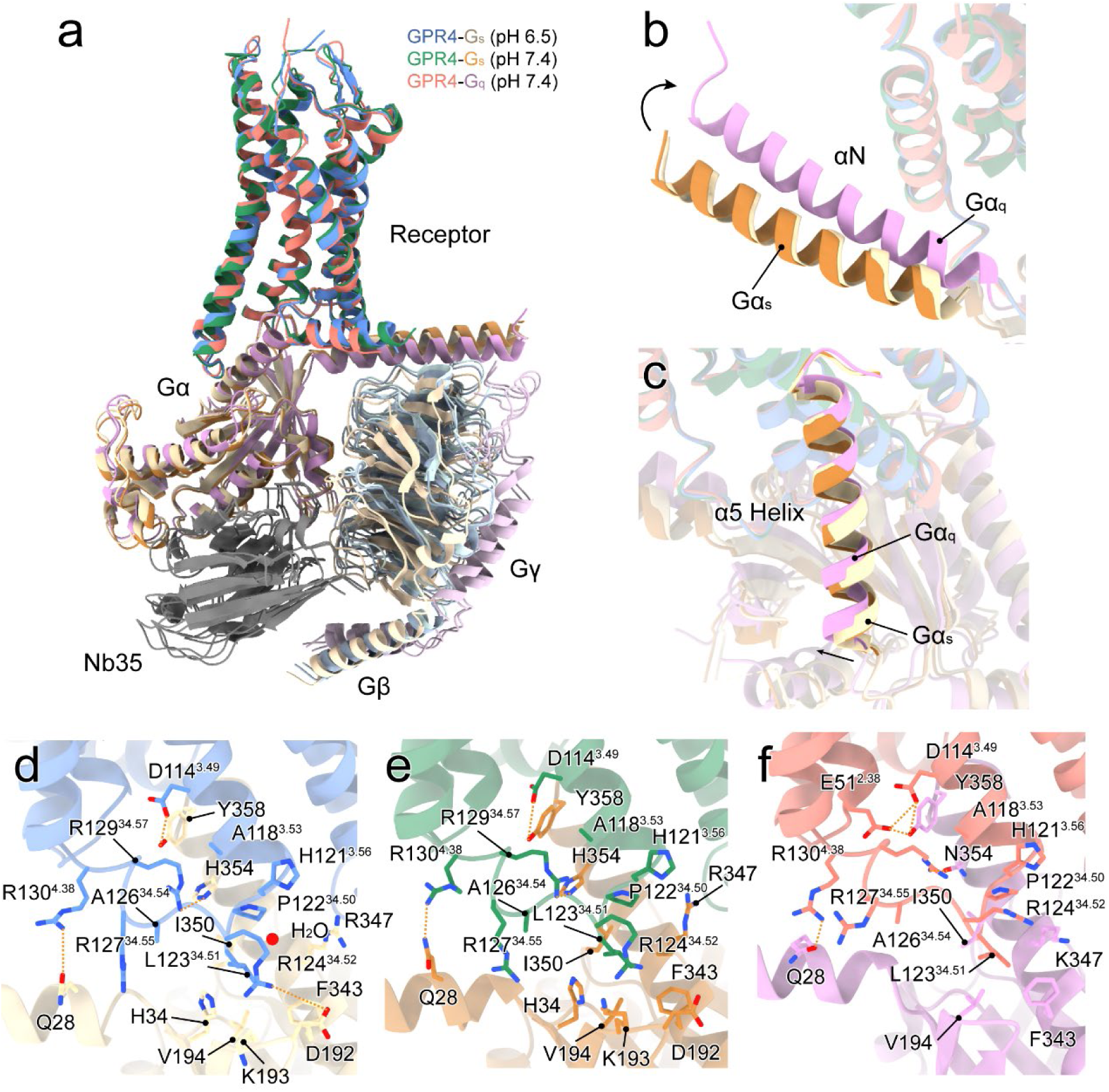
G protein interfaces between active GPR4 and Gα_s_/Gα_q_. **a** Overall structural comparison of GPR4 complexes. **b-c** Movement of αN and α5 Helix of Gα_s_/Gα_q_. The directions are shown by black arrows. **d-f** Detail interactions between GPR4 and Gα subunits. Interactions between GPR4 at different pH conditions and Gα_s_ are displayed in **d** (pH 6.5) and **e** (pH 7.4), respectively. Interactions between GPR4 at pH 7.4 and Gα_q_ are displayed in **f**. Colors are shown as indicated.

**Fig. S13.**
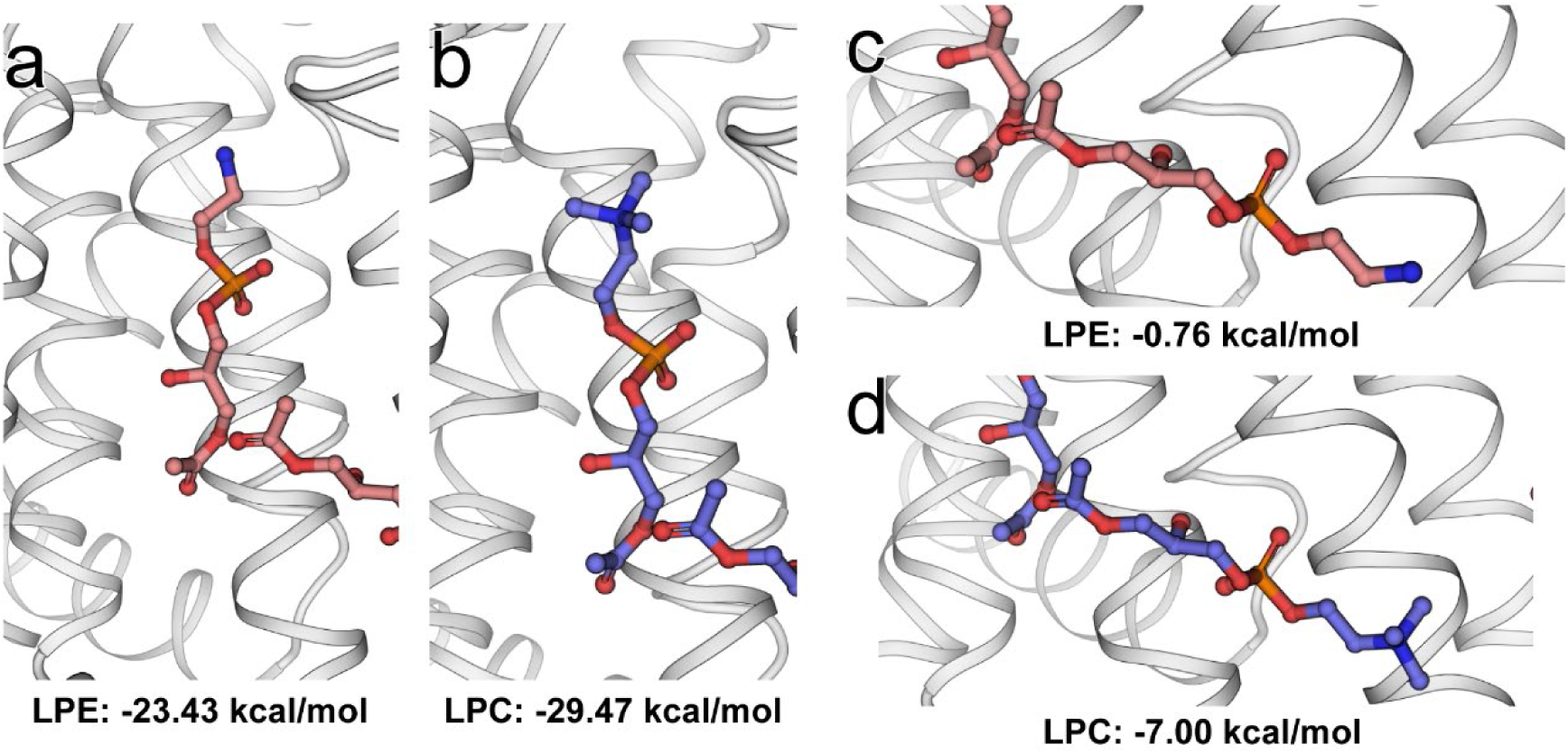
Binding free energy estimation of highly similar LPC analogues in its pocket at the top of intracellular TM4-TM5 (a-b), and at the bottom of intracellular TM4-TM5 (c-d). The binding free energy value is shown below.

**Table S1.** Cryo-EM data collection, refinement and validation statistics.

|  | GPR4-G<sub>s</sub> (pH<br>6.5)<br>(EMD-61370)<br>(PDB: 9JCO) | GPR4-G<sub>s</sub> (pH<br>7.4)<br>(EMD-61372)<br>(PDB:9JCQ) | GPR4-G<sub>q</sub> (pH<br>7.4)<br>(EMD-61371)<br>(PDB: 9JCP) |
| --- | --- | --- | --- |
| **Data collection and processing** |  |  |  |
| Magnification | 105,000 | 64,000 | 105,000 |
| Voltage (kV) | 300 | 300 | 300 |
| Electron exposure (e<sup>-</sup>/Å<sup>2</sup>) | 50 | 50 | 50 |
| Defocus range (μm) | -1.0~-3.0 | -1.0~-3.0 | -1.0~-3.0 |
| Pixel size (Å) | 0.73 | 0.824 | 0.73 |
| Symmetry imposed | C1 | C1 | C1 |
| Initial particle images (no.) | 2,798,739 | 3,349,414 | 4,555,181 |
| Final particle images (no.) | 596,409 | 626,376 | 550,055 |
| Map resolution (Å) | 2.36 | 2.59 | 2.55 |
| FSC threshold | 0.143 | 0.143 | 0.143 |
| Map resolution range (Å) | 2-4 | 2-4 | 2-4 |
| **Refinement** |  |  |  |
| Initial model used (PDB code) |  |  |  |
| Model resolution (Å) | 2.2 | 2.4 | 2.6 |
| FSC threshold | 0.143 | 0.143 | 0.143 |
| Model resolution range (Å) | 50-2.2 | 50-2.4 | 50-2.6 |
| Map sharpening *B* factor (Å<sup>2</sup>) | -50 | -60 | -60 |
| Model composition |  |  |  |
| Non-hydrogen atoms | 8431 | 8437 | 8332 |
| Protein residues | 1051 | 1052 | 1048 |
| Ligands | 2 | - | - |
| *B* factors (Å<sup>2</sup>) |  |  |  |
| Protein | 54.58 | 46.49 | 66.39 |
| Ligand | 58.15 | 41.14 | 47.25 |
| Water | 76.42 | 53.88 | 61.67 |
| R.m.s. deviations |  |  |  |
| Bond lengths (Å) | 0.006 | 0.006 | 0.004 |
| Bond angles (°) | 1.034 | 0.723 | 0.617 |
| Validation |  |  |  |
| MolProbity score | 1.55 | 1.31 | 1.30 |
| Clashscore | 7.06 | 5.31 | 3.94 |
| Poor rotamers (%) | 1.00 | 0.78 | 0.79 |
| Ramachandran plot |  |  |  |
| Favored (%) | 97.11 | 97.88 | 97.39 |
| Allowed (%) | 2.89 | 2.12 | 2.61 |
| Disallowed (%) | 0.00 | 0.00 | 0.00 |

